# Modulation of motor vigour by expectation of reward probability trial-by-trial is preserved in healthy ageing and Parkinson’s disease patients

**DOI:** 10.1101/2022.08.25.505068

**Authors:** Margherita Tecilla, Michael Grossbach, Giovanni Gentile, Peter Holland, Angelo Antonini, Maria Herrojo Ruiz

## Abstract

Motor improvements, such as faster movement times or increased velocity, have been consistently associated with reward magnitude in deterministic contexts. Yet whether individual inferences on reward probability influence motor vigour dynamically remains undetermined. Here we investigated how dynamically inferring volatile action-reward contingencies modulated motor performance trial-by-trial in healthy younger (HYA, 37) and older adults (HOA, 37), and in medicated Parkinson’s Disease patients (PD, 20). We conducted an online study that coupled a standard one-armed bandit decision-making paradigm with a motor sequence task and used a validated hierarchical Bayesian model to fit trial-by-trial data. Our results showed that stronger predictions about the tendency of the action-reward contingency led to faster performance tempo on a trial-by-trial basis without modulating reaction times (RT). Using Bayesian linear mixed models, we demonstrated in HYA, HOA and PD a similar sensitivity (slope) of execution tempo to inferences about the reward probabilities, despite HOA and PD being generally slower than HYA (intercept). In a second experiment in HYA (39), we additionally showed that subjective inferences about credit assignment – whether lack of reward is associated with an incorrect decision or execution error – led to a similar modulation of motor vigour by reward expectation. Our study is the first to reveal that the dynamic updating of beliefs about volatile action-reward contingencies positively biases motor performance through faster execution tempo, without affecting RT. We also provide novel evidence for a preserved sensitivity of motor vigour to inferences about the action-reward mapping in ageing and medicated PD.

**SIGNIFICANCE STATEMENT:** Navigating a world rich in uncertainty relies on updating beliefs about the probability that our actions lead to reward. Here we investigated how inferring the action-reward contingencies in a volatile environment modulated motor vigour trial-by-trial in healthy younger and older adults, and in Parkinson’s Disease patients on medication. We found an association between trial-by-trial predictions about the tendency of the action-reward contingency and performance tempo, with stronger expectations speeding performance. We additionally provided evidence for a similar sensitivity of performance tempo to the strength of these predictions in all groups. Thus, dynamic beliefs about the changing relationship between actions and their outcome invigorated motor performance. This positive bias was not compromised by age or Parkinson’s disease.

## INTRODUCTION

The prospect of obtaining rewards invigorates motor performance, with incentives leading to faster and more accurate movements (Summerside et al., 2018; Sedaghat-Nejad et al., 2019; Codol et al., 2020). Several non-mutually exclusive mechanisms have been proposed to account for the beneficial effects of reward on movement. These include the reward-driven strengthening of motor representations at the cortical level (Galaro et al., 2019; Adkins & Lee, 2021), enhanced feedback-control processes (Padmala & Pessoa, 2011; Botvinick & Braver, 2015; Carroll et al., 2019; Manohar et al., 2019), increased limb stiffness (Codol et al., 2020) and coarticulation (Sporn et al., 2020; Aves et al., 2021). Despite the growing number of studies demonstrating how rewards positively bias motor behaviour, the evidence so far is limited to simple manipulations of reward magnitude (presence/absence; large/small). Yet, in our everyday life we are exposed to environments rich in uncertainty, where adaptive behaviour relies on estimating the changing relationship between actions and their outcomes. How beliefs about the probabilistic structure of reward contingencies modulate motor performance remains largely unexplored. In addition, whether this modulation is compromised with age and in neurological conditions is unclear.

Hierarchical Bayesian inference models explain how individuals learn and make decisions under uncertainty (Behrens et al., 2007; den Ouden et al., 2010; Feldman & Friston, 2010). In multi/one-armed bandit tasks, these models describe learning as governed by inferences on the probabilistic stimulus-outcome mappings, as well as higher-level beliefs about the rate of change of these contingencies over time, labelled volatility (Friston et al., 2014; Mathys et al., 2014; de Berker et al., 2016; Sheffield et al., 2022). In Bayesian predictive coding, beliefs about the probable causes of sensory data are updated via prediction errors weighted by uncertainty or precision (Friston et al., 2014; Mathys et al., 2014). Thus, dynamic estimates of uncertainty allow for the expression of individual differences in belief updating. If motor vigour is modulated by beliefs about the action-reward contingencies, then individual differences in uncertainty estimates could explain differences in motor vigour. Alternatively, under equivalent signatures of decision-making behaviour, individuals could exhibit differential sensitivity of motor performance to the expectation of reward probability.

Here we tested these hypotheses by determining whether dynamic predictions about volatile action-reward contingencies influence motor sequence performance trial-by-trial. We additionally assessed whether the sensitivity of motor performance to the strength of these predictions undergoes changes in later stages of life and in patients with Parkinson’s Disease (PD) on their dopamine-replacement medication. This is motivated by the lack of evidence regarding how reward sensitivity and reversal learning interact to modulate motor vigour in PD and healthy ageing. On the one hand, evidence supports preserved sensitivity to rewards and probabilistic learning in healthy ageing and medicated PD (Fera et al., 2005; Euteneuer et al., 2009; Pietschmann et al., 2011; Gescheidt et al., 2012, 2013; Chen et al., 2020; Aves et al., 2021). Yet other work suggests impoverished decision making and reward-based learning in both groups. Specifically, ageing and medicated PD can underperform in tasks using volatile probabilistic stimulus-outcome mappings (Cools et al., 2001; Shohamy et al., 2004; Weiler et al., 2008 Eppinger et al., 2011; Nassar et al., 2016; Hämmerer et al., 2019). However, the medication effects on decision making in PD (on/off states) is still under debate (Ryterska et al., 2013; Kjær et al., 2019). Accordingly, whether ageing and medicated PD can use their dynamic belief estimates to invigorate motor performance trial-by-trial remains unspecified.

To address our questions, we conducted two online experiments that used a reward-based motor decision-making task based on a one-armed bandit paradigm with changing stimulus-outcome contingencies over time. In the first study we assessed the influence of the strength of predictions about the action-reward contingency on motor performance on a trial-by-trial basis. We also investigated whether this modulation is compromised in ageing and PD. In the second control study we evaluated the potential contribution of subjective inferences about credit assignment to explain our results.

## MATERIALS AND METHODS

### Participants

37 healthy younger adults (HYA; 13 males, age 18-40, mean age 27.8, standard error of the mean [SEM] 0.67; hereafter we follow the intrinsic measures of precision for rounding descriptive and inferential statistics as reported in Cousineau (2020), 20 patients diagnosed with Parkinson’s Disease (PD; 13 males, age 40-75, mean age 58.9, SEM 1.32) and an age-matched group of 37 healthy older adults (HOA; 20 males, age 40-75, mean age 61.5, SEM 1.25) participated in this research. The sample size for healthy samples was informed by previous work assessing differences between HYA and HOA in decision-making under uncertainty (de Boer et al., 2017: N = 30, 30) and our own work assessing group effects in parameters of hierarchical Bayesian models (Hein et al., 2021; 2022; N = 20, 20). We increased the sample size to allow for variability being introduced due to the nature of the online study.

The study received ethical approval by the review board of Goldsmiths (healthy sample), University of London, and the Neurology Clinic, Padua University Hospital (PD sample). Informed consent was acquired for each participant. HYA and HOA were recruited through online advertisement and via the Research Participation Scheme (RPS) at Goldsmiths University, while PD were enrolled at the Neurology Clinic, Padua University Hospital. All participants were right-handed, had normal or corrected vision and were able to perform controlled finger movements. Amateur/professional pianists and participants diagnosed with a mental health disorder were excluded from the study. Additionally, exclusion criteria for PD patients were: implanted with Deep Brain Stimulation (DBS), taking antidepressant medications, diagnosed with dementia and displaying tremor as an onset symptom. One PD patient declared to take Laroxyl, yet confirmed not to be diagnosed with depression. PD were evaluated through ITEL-Mini Mental state examination (ITEL-MMSE; Metitieri et al., 2001), Unified Parkinson’s Disease Rating Scale part III (UPDRS-III; Fahn & Elton, 1987), Hospital Anxiety and Depression Scale (HADS; Zigmond & Snaith, 1983) and State-Trait Anxiety Inventory (STAI Y2; Spielberger, 1983). Supplementary disease-related information was also gathered (**Table 1**). Patients completed the experiment in the ON medication state according to their usual dopamine-replacement treatment. The individual dopaminergic medication details were collected and converted to a levodopa-equivalent daily dose (LEDD) value (**Table 1**).

**Table 1.**
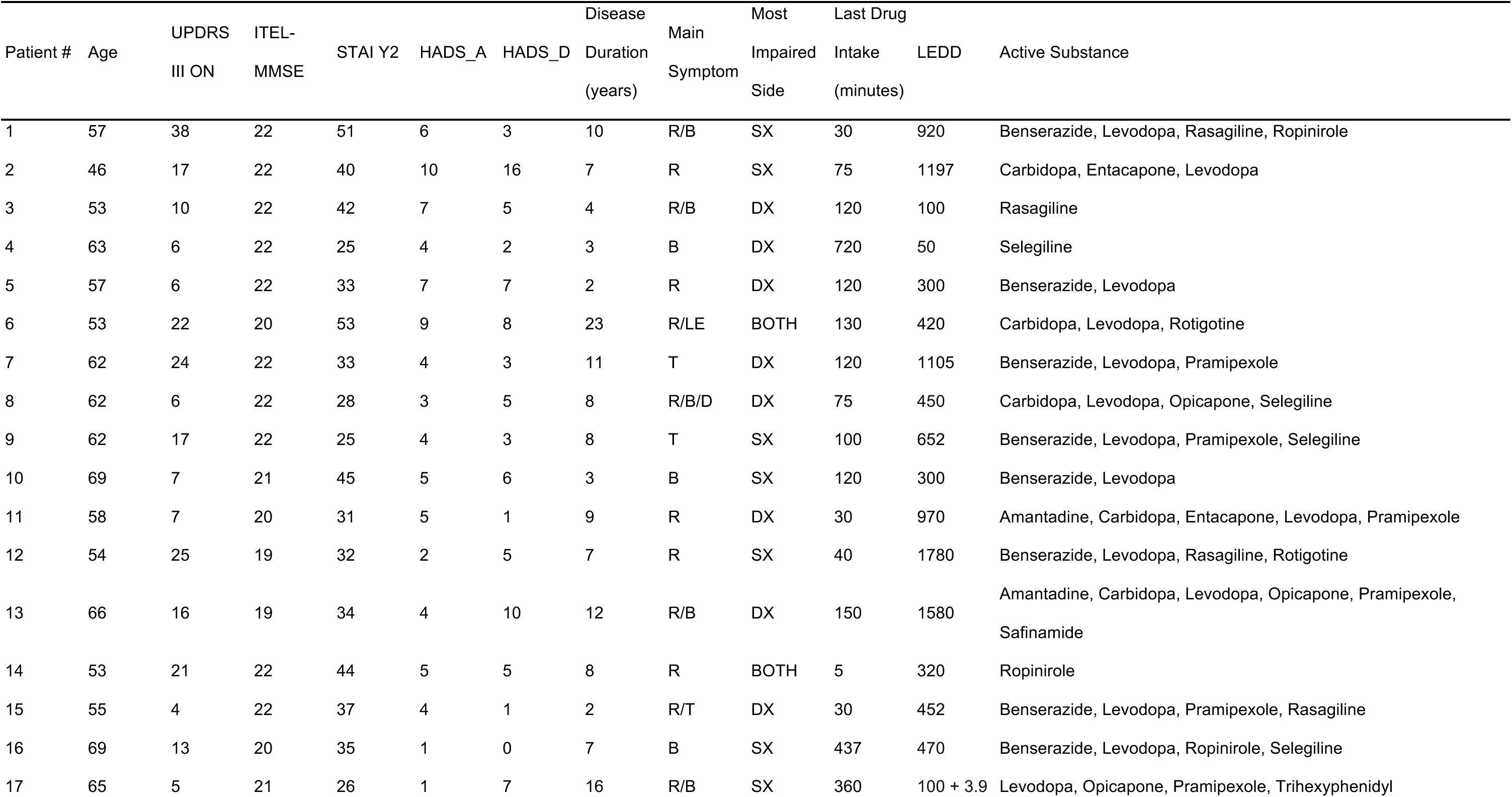
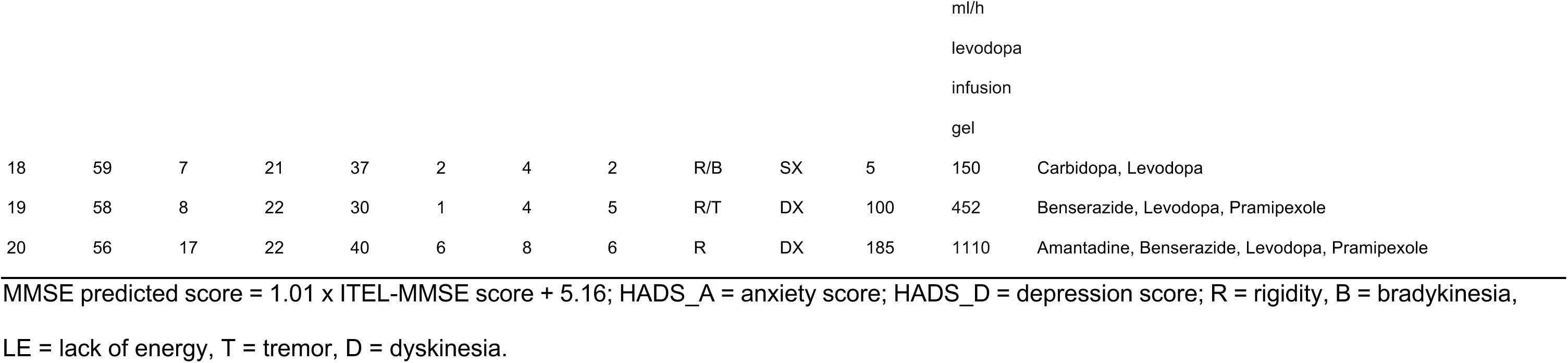
PD clinical information.

All participants took part in the study remotely (online), except for five PD patients, who completed the study in the laboratory facilities of the Neurology Clinic of Padua. An Italian translation of the original experimental instructions in English was created to test some of the HOA participants (N = 24) and all PD patients (see the Results section for details on our control analyses to assess the effect of the language of the instructions). The previously validated Italian translations of the HADS, ITEL-MMSE, UDPRS-III and STAI Y2 scales were used. HYA and HOA participants received a monetary compensation of £5 (5€ for those completing the task in Italian), which could be increased up to £10 (10€) as a function of their task performance. PD patients did not receive a monetary prize, in line with the clinical research policies at the Neurology Clinic of Padua.

A separate sample of 39 HYA took part in a second control study aimed at evaluating the potential contribution of subjective inferences about task-related reward (credit) assignment to explain our results (McDougle et al., 2016). The same recruitment strategies, inclusion/exclusion criteria and compensation as for HYA in the main experiment applied. HYA participants in this control experiment were divided into two subsamples as a function of their reply (True/False) to a post-performance question (Q8; **Table 2**). Group Q8_T_ consisted of 26 participants (8 males, age 18-40, mean age 24.1, SEM 1.13) and Q8_F_ of 13 participants (2 males, age 18-40, mean age 25, SEM 1.7).

**Table 2.**
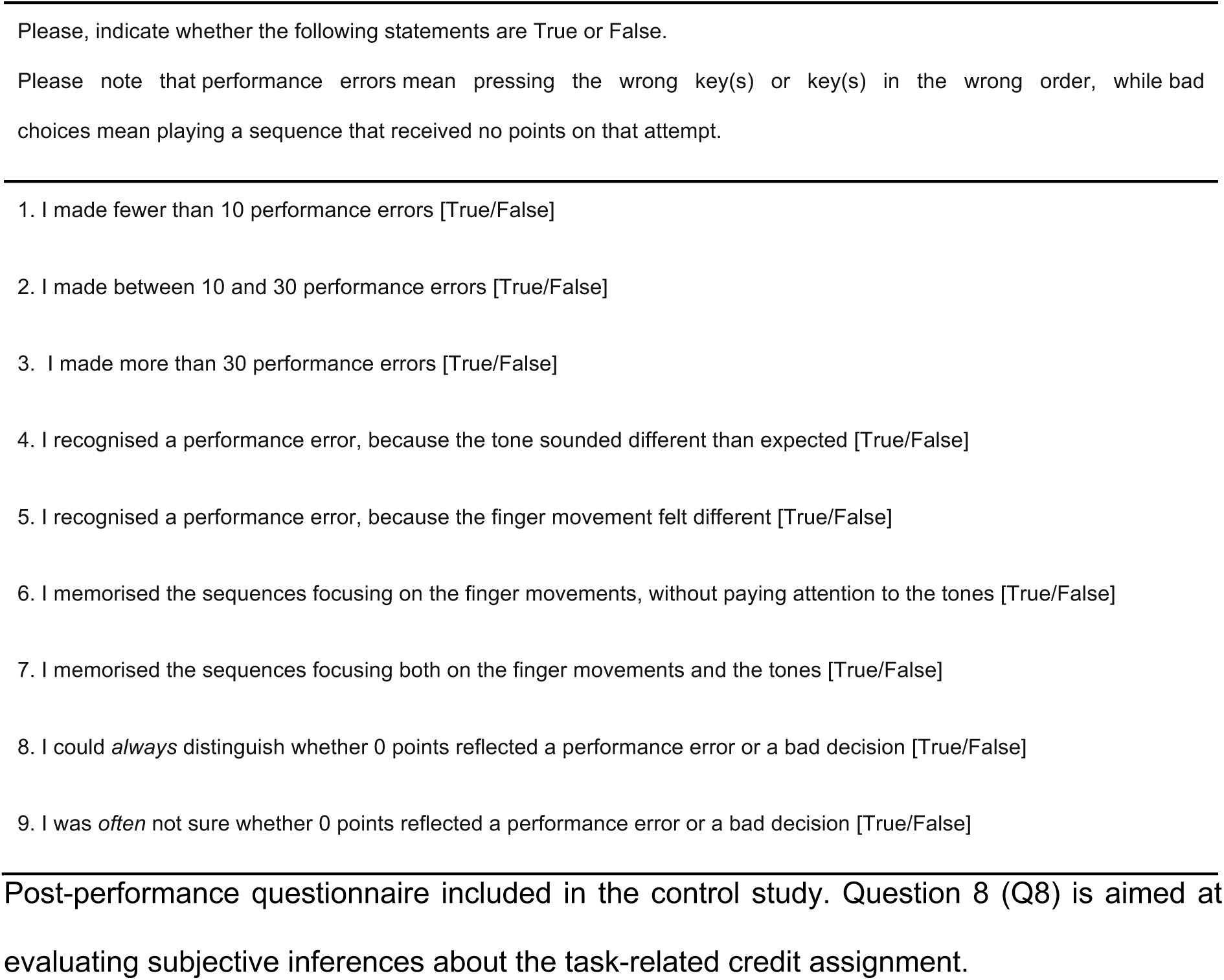
Post-performance questionnaire.

### Experimental design

The experiment ran completely online on the Qualtrics platform (https://www.qualtrics.com) and was accessible through a study link. The task was programmed in JavaScript and embedded into the Qualtrics form. We provide more details of the data acquisition below (see Acquisition of online data using JavaScript section).

Participants performed a novel computerised reward-based motor decision-making task based on a one-armed bandit paradigm with changing stimulus-outcome contingencies over time (e.g., de Berker et al., 2016). Participants were instructed to play one of two sequences of finger movements on a virtual piano to express their decision, which is an extension of standard one-armed bandit tasks that instruct participants to manifest their choice by pressing a right or left button (Hein et al., 2021).

The task consisted of a familiarisation and a reward-based learning phase. In the familiarisation phase participants learned how to play two short sequences (seq1 and seq2) of four finger presses each. Each sequence was uniquely represented by one of two different fractal images (**Figure 1A**). They were asked to position their right hand on the keyboard as follows: index finger on “g” key, middle finger on “h” key, ring finger on “j” key and little finger on “k” key. Each key press reproduced a distinct auditory tone, simulating a virtual piano. Participants were trained to press “g-j-h-k” for seq1 (red fractal) and “k-g-j-h” for seq2 (blue fractal). Online videos showing the correct hand position on the keyboard and how to perform the two sequences were provided to increase inter-individual consistency. The familiarisation phase terminated when an error-free performance was achieved for five times in succession for both sequences. The number of sequence renditions during familiarisation was recorded and used for subsequent analyses.

**Figure 1.**
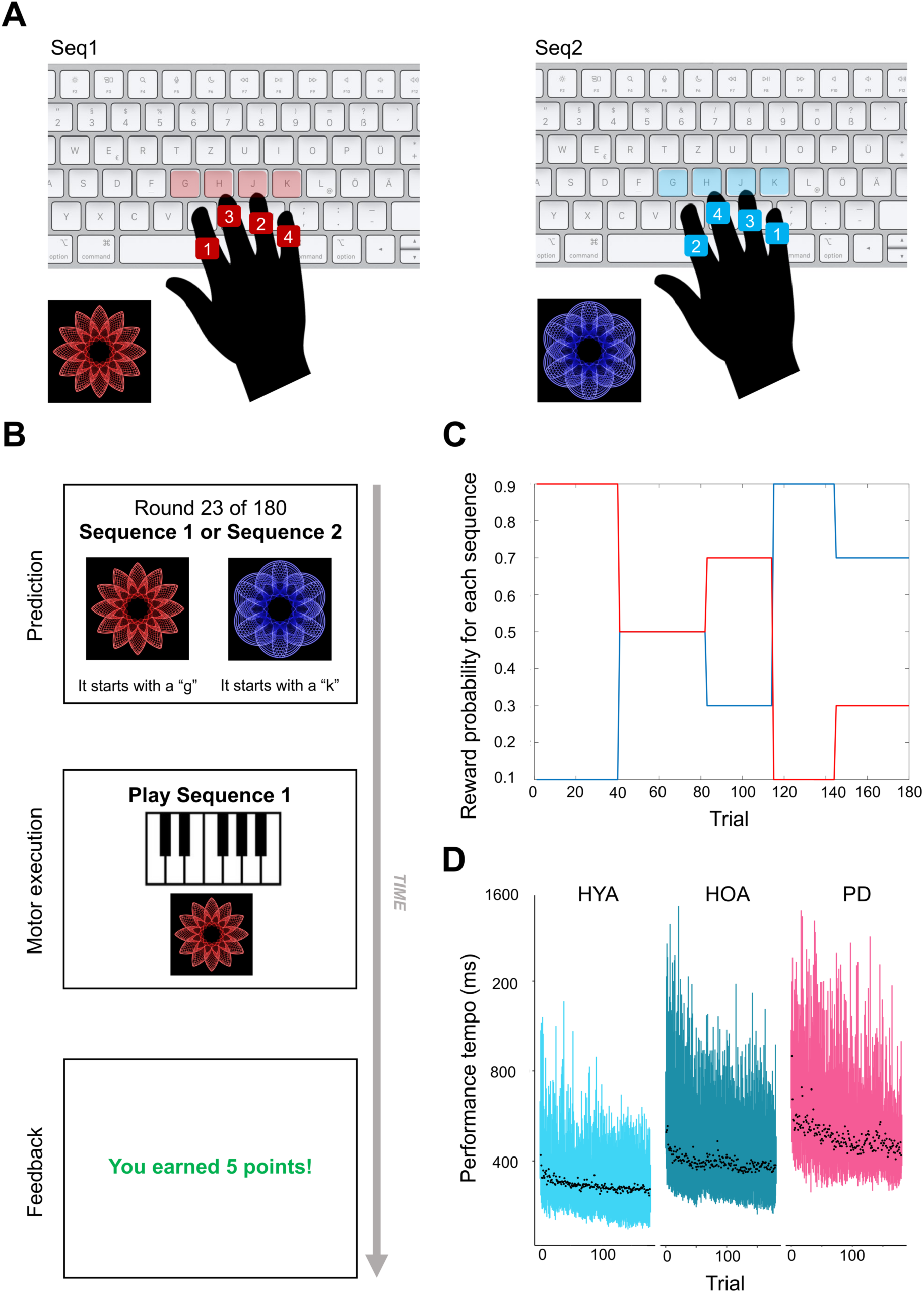
Task structure. ***A***, In the task familiarisation phase participants learnt to play two sequences associated with two images (red fractal – seq1 “g-j-h-k”; blue fractal – seq2 “k-g-j-h”). ***B***, On each trial of the reward-based learning phase, subjects decided which sequence to play in order to get the reward. The two icons were always either red or blue and presented to the left or right part of the screen, respectively. First, participants made a prediction about which sequence (associated to the corresponding icon) was more likely to give them a reward. When a decision was reached, they played the corresponding sequence using the keyboard. Finally, the outcome (win +5p or 0p) was revealed. In the example, the participant played seq1 and obtained five points, suggesting correct prediction and execution. ***C***, Displays the typical subject-specific mapping of probabilistic stimulus-outcome contingency over the course of 180 trials. In the example, the order of reward mappings for the blue icon (and corresponding seq2) is 10-50-30-90-70% (reciprocal for red icon and corresponding seq1). In order to obtain the maximal reward, participants needed to track these changes and adapt their choices throughout the experiment. ***D***, The trial by trial changes in performance tempo in ms (mIKI; mean inter-keystroke-intervals; see Behavioural and computational data analysis section for further details) for healthy younger adults (HYA; light blue), healthy older adults (HOA; dark blue) and patients with Parkinson’s Disease (PD; in purple) across 180 trials. Black dots represent the trial-by-trial within-group averages of performance tempo. Bars indicate 95% interval probabilities. Participants tended to play the sequences faster towards the end of the experiment, possibly reflecting a practice effect.

The reward-based learning phase consisted of 180 trials. On each trial, participants were instructed to choose between two coloured fractals (blue and red) and correctly play the associated sequence (seq1 and seq2) in order to receive a reward (five points; **Figure 1B**). Trial-by-trial reward feedback about participants’ choices was provided on the screen (binary: “You earned 5 points!” or “You earned 0 points”). The reward probability associated with each sequence (or icon) changed every 25-35 trials (as in de Berker et al., 2016). The mapping governing the likelihood of sequences being rewarded was reciprocal (p(win|seq1) = 1- p(win|seq2)) and consisted of five stimulus-outcome contingency blocks (90/10, 70/30, 50/50, 30/70, 10/90) (**Figure 1C**). The order of the contingency blocks was randomly generated for each participant.

The sequence performance had to be completed within 5000 ms, terminating in a Stop signal. Visual hints suggesting the first key to press for both sequences were displayed: “It starts with a “g”” – for seq1 (red fractal); “It starts with a “k”” – for seq2 (blue fractal). Participants were instructed to press key “q” if they needed a reminder of the order of finger presses for each sequence. No participant required this reminder.

Correctly playing the rewarded sequence added five points to the participants’ total score (win trial). Thus, receiving five points indicated that participants chose the rewarded sequence on the trial and did not make performance execution errors when playing it. Zero points, however, could reflect participants choosing an unrewarded sequence on that trial or, alternatively, choosing a rewarded sequence but performing it incorrectly (performance execution error) (McDougle et al., 2016). No reward was provided when sequence performance exceeded the 5000 ms limit (no response trial) and participants were informed they played too slowly.

Thus, to maximise the total cumulative points over the experiment, participants had to infer the probability of reward associated with each sequence and adapt their choices when contingencies changed. They also had to perform the sequences correctly. Participants were informed at the beginning of the experiment that the stimulus-outcome mapping would change from time to time. However, they received no detailed information regarding the frequency or magnitude of those changes. We validated that all participants completed the task correctly by assessing the percentage of trials that they performed either seq1 or seq2. If participants’ decision to play one of the sequences was as a function of the inferred tendency of the stimulus-outcome contingency, they should play each sequence in 50% of the trials on average. The reverse, however, was not necessarily true. See further details below (Behavioural and computational data analysis and Results sections).

The control study followed the same procedure as the main experiment. The only difference consisted of asking participants at the end of the reward-based learning phase to reply to some questions about their performance. We were particularly interested in assessing whether participants could correctly infer what zero points meant, that is, whether they could distinguish between a performance execution error or a decision to play a sequence that was unrewarded on the trial. Both scenarios would result in zero points. **Table 2** lists the questions of the post- performance questionnaire, which required binary responses (True/False) and was designed based on previous work (McDougle et al., 2016; Herrojo Ruiz et al., 2017). The binary answer to Question 8 “I could *always* distinguish whether 0 points reflected a performance error or a bad decision” was used as criterion to split the control sample into Q8_T_ (i.e., participants were *always* sure about the hidden causes for the lack of reward) and Q8_F_ (i.e., participants were *not always* sure about the hidden causes for receiving zero points). Among other questions, participants were asked whether the subjective number estimate of performance errors was less than 10, between 10 and 30 or more than 30. This information was used to investigate whether Q8_T_ and Q8_F_ differed in the rate of subjective execution errors. The rationale here was that Q8_F_ participants relative to Q8_T_ could attribute more zeros to performance errors rather than inferring that their choice was not rewarded on that trial. Alternatively, they could misattribute zeros to bad decision outcomes. In both cases, their biased credit assignment would be reflected in a more pronounced difference between estimated and empirical error rates in Q8_F_. However, their belief updating would differ; in the first case, Q8_F_ participants relative to Q8_T_ would not update their beliefs following a zero outcome, as this would be rendered as not informative feedback regarding the underlying probabilistic structure. Thus, differences in credit assignment could explain differences in decision making and, potentially, associated motor vigour effects.

Finally, we also assessed the strategy that participants used to memorise the sequences (79.5% of participants declared to have memorised the sequences focusing both on the finger movements and the tones; Q7).

### Acquisition of online data using JavaScript

Due to the nature of the online experiment, cross-browser issues could emerge. A potential issue was that participants could use a variety of computer hardware, running on different web browsers operating systems and keyboard types (e.g., tablets vs laptops). To mitigate the effect of hardware variability on the acquisition of motor performance data, we instructed participants to complete the task on a desktop or laptop computer. An inspection of browser user agent data suggests that the experiment was performed on a mixture of desktops or laptops running the Chrome & Safari browsers on Windows and Macintosh operating systems. Timing data was collected using the web browser’s high-resolution timer. This browser resolution timer has an upper resolution limit of 2 ms on some web browsers. Therefore, all analysis scripts *truncated* timing data to 2 ms precision. When estimating the mean and standard error of the mean in time variables, we therefore considered a systematic error of 1 ms (2 ms precision means that our time measures were on average 1 ms too short). For each participant, keypresses, timing data, points, contingency mapping, outcome, and other data were extracted on each trial, then stored and uploaded via JSON to the data folder in Pavlovia (see https://gitlab.pavlovia.org/oshah001/reward-learning-experiment).

### The hierarchical gaussian filter

To model intra-subject trial-by-trial performance in our task, we used a validated hierarchical Bayesian inference model, the Hierarchical Gaussian Filter (HGF; Mathys et al. 2011, 2014; Frässle et al., 2021). The HGF toolbox is an open source software and is freely available as part of TAPAS (http://www.translationalneuromodeling.org/tapas; Frässle et al., 2021). Here we used the HGF version 6.1 implemented in MATLAB® 2020b. The HGF is a generative model that describes how individual agents learn about a hierarchy of hidden states in the environment, such as the latent causes of sensory inputs, probabilistic contingencies, and their changes over time (labelled volatility). Beliefs on each hierarchical level are updated through prediction errors (PEs) and scaled (weighted) by a precision ratio (precision as inverse variance or uncertainty). The precision ratio effectively operates as a learning rate, determining how much influence the uncertainty about the belief distributions has on the updating process (Mathys et al., 2011, 2014).

In the present study, the HGF was used to characterise subject-specific trial-by-trial trajectories of beliefs about stimulus-outcome contingencies (level 2) and their changes over time (environmental volatility, level 3). These belief distributions are Gaussian, summarised by the posterior mean (*µ*_2_, *µ*_3_) and the posterior variance (*σ*_2_, *σ*_3_). The latter represents uncertainty about the hidden states on those levels, that is, our imperfect knowledge about the true hidden states. On level 2, *σ*_2_ is termed estimation or informational uncertainty. More generally, the inverse 1/*σ* is termed precision, labelled π. The HGF provides trajectories of updated beliefs on the current trial, *k*, after observing the outcome (*µ*_i_^(k)^ for level *i* = 2, 3), and therefore also predictions about those states on the previous trial (*µ*_i_^(k-1)^, also denoted by the hat operator *µ_i_^(k)^*).

As in previous work using one-armed bandit paradigms (Iglesias et al., 2013; Mathys et al., 2014; Hein et al., 2021), we modelled learning using the 3-level HGF (HGF_3_) for binary outcomes (**Figure 2A**). In this hierarchical perceptual model, the hidden state on the lowest level, x_1_, represents the binary categorical variable of the experimental stimuli (for each trial *k*, x_1_^(k)^ = 0 if the red icon/seq1 is rewarded [or blue/seq2 loses]; x_1_^(k)^ = 1 when red fractal/seq1 is not rewarded [or blue/seq2 wins]). Higher in the hierarchy, x_2_ reflects the true value of the tendency of the stimulus-outcome contingency, and x_3_ the true volatility of the environment (i.e., of x_2_). Belief updating in the HGF depends on various parameters, which can be estimated in each individual or fixed depending on the hypotheses. This allows for the assessment of individual learning characteristics. Here we chose to individually estimate parameter ω_2_, representing the tonic (time-invariant) volatility on the second level, and ω_3_, denoting the tonic volatility on the third level. Generally, ω_2_ and ω_3_ parameters describe an individual’s learning motif. Larger ω_2_ values are associated with faster learning about stimulus outcomes, and thus greater update steps in *µ*_2_ (see simulations in Hein et al., 2021). Similarly, greater levels of tonic volatility on level 3, ω_3_, increase the update steps on *µ*_3_. See details onour priors in **Table 3.** Using simulations to assess the accuracy of parameter estimation in the HGF_3_, we and others have previously demonstrated that ω_2_ can be estimated accurately, while ω_3_ is not estimated well (Reed et al., 2020; Hein et al., 2021).

**Figure 2.**
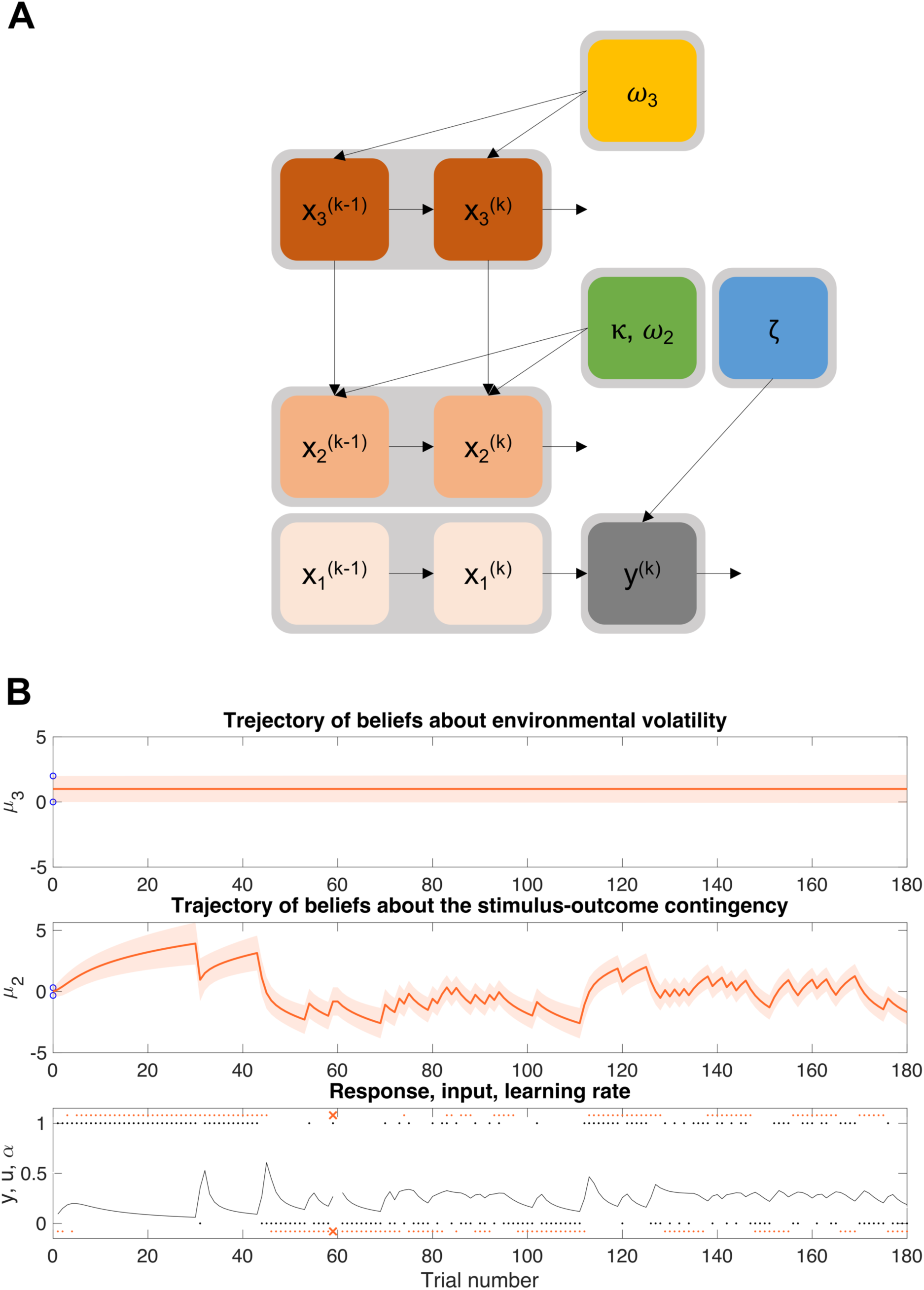
The Hierarchical Gaussian Filter (HGF) for binary outcomes. ***A,*** Illustration of the 3- level HGF model (HGF_3_) with relevant parameters modulating each level (adapted from Hein et al., 2021). Level *x_1_* represents the binary categorical variable of the experimental stimuli on each trial *k*; *x*_2_ reflects the true value of the tendency of the stimulus-outcome contingency, and *x*_3_ the true volatility of the environment. In our experiment, *ω*_2_, *ω*_3_ and, were free parameters and were estimated by fitting individual responses and observed inputs with the HGF. - represents the strength of coupling between level 2 and 3 (fixed to 1 in our study; not shown in the text; see Mathys et al., 2014 for the model equations). ***B,*** Belief trajectories for the HGF_3_ across the total 180 trials in a representative participant. At the lowest level, black dots (*u*) represent the outcomes, denoting whether seq1 was rewarded or notl (1 = seq1 wins [seq2 loses]; 0 = seq2 wins [seq1 loses]); orange dots (*y*) represent the participant’s choices (1 = seq1 is played; 0 = seq2 is played); orange crosses depict performance execution errors; the black line is a subject-specific learning rate about stimulus outcomes (.; see Mathys et al. 2014 for the full HGF equations). At the second level, *µ*_2_ (*σ*_2_) is the trial-by-trial trajectory of beliefs (mean and variance) about the tendency of the stimulus-outcome contingencies (*x*_2_). A mean estimate *µ*_2_ shifted towards positive values on the y-axis indicates that the participant had a greater expectation that seq1 was rewarded relative to seq2. In addition, larger (absolute) *µ*_2_ values on that axis denote a stronger expectation that given the correct sequence choice a reward will be received. The trajectory of beliefs about phasic (log)volatility (*µ*_3_ [*σ*_3_]) is displayed at the top level. The true volatility in our task, *x*_3_, was constant, as the stimulus-outcome contingencies changed every 25-35 trials. Participants could, however, express individual differences in their log-volatility estimates, which could be captured by the HGF_3_ (e.g., Powers et al., 2017). In our study, the winning model was the 2-level HGF (HGF_2_), in which volatility was fixed across participants. Blue circles on the y-axis denote the upper and lower priors of the posterior distribution of beliefs, *µ*_i_^(0)^± “_i_^(0)^, *i* = 2,3.

**Table 3.**
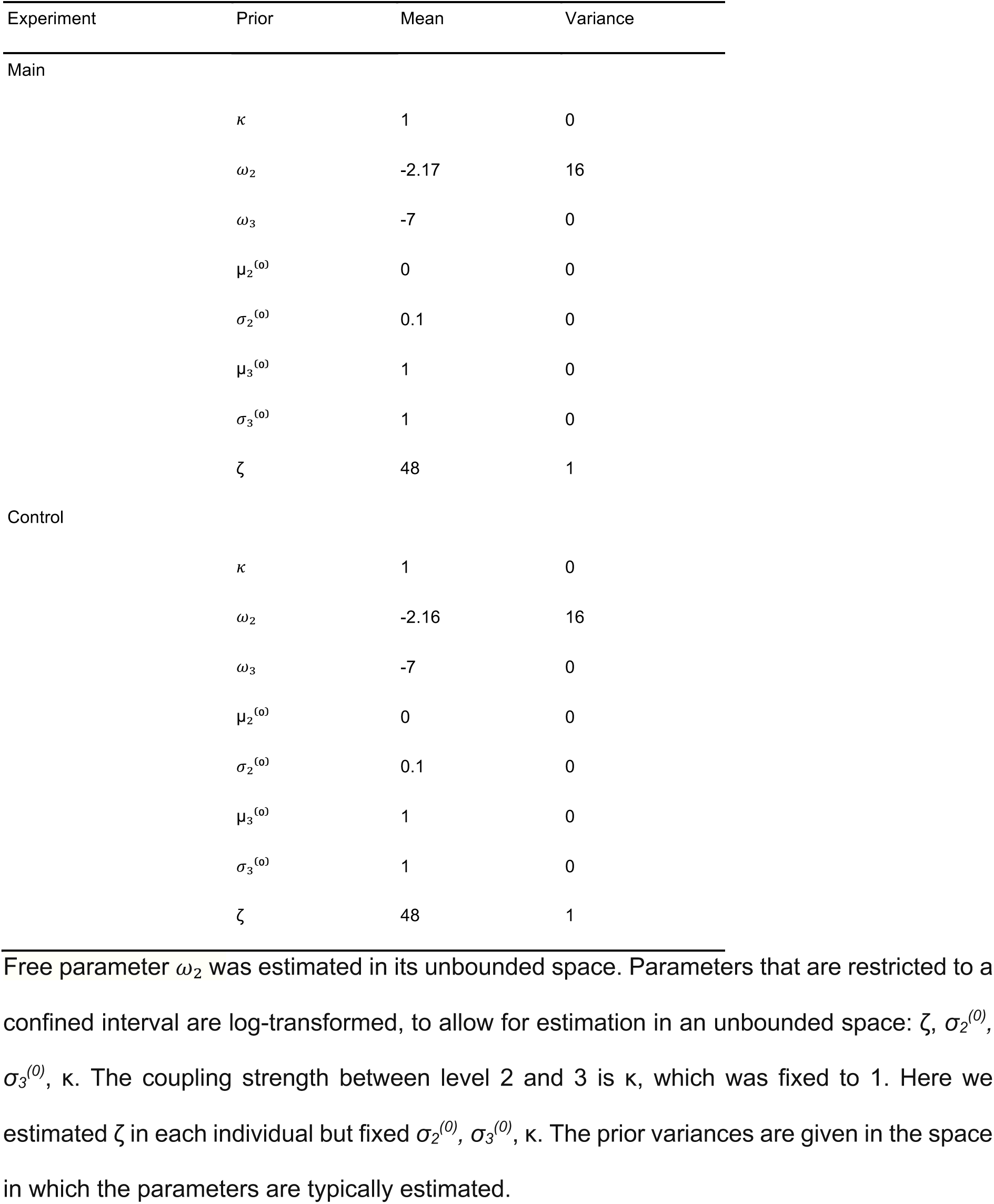
Means and variances of the priors on perceptual parameters and starting values of the beliefs of the winning HGF model for the main and control experiments.

| Experiment | Prior | Mean | Variance |
| --- | --- | --- | --- |
| Main |  |  |  |
| | $\kappa$ | 1 | 0 |
| | $\omega_2$ | -2.17 | 16 |
| | $\omega_3$ | -7 | 0 |
| | $\mu_2^{(0)}$ | 0 | 0 |
| | $\sigma_2^{(0)}$ | 0.1 | 0 |
| | $\mu_3^{(0)}$ | 1 | 0 |
| | $\sigma_3^{(0)}$ | 1 | 0 |
| | $\zeta$ | 48 | 1 |
| Control |  |  |  |
| | $\kappa$ | 1 | 0 |
| | $\omega_2$ | -2.16 | 16 |
| | $\omega_3$ | -7 | 0 |
| | $\mu_2^{(0)}$ | 0 | 0 |
| | $\sigma_2^{(0)}$ | 0.1 | 0 |
| | $\mu_3^{(0)}$ | 1 | 0 |
| | $\sigma_3^{(0)}$ | 1 | 0 |
| | $\zeta$ | 48 | 1 |

We then coupled the perceptual HGF model to a response model for binary outcomes, which defined how beliefs about the tendency of the stimulus-outcome contingencies were mapped onto decisions (e.g., which sequence should be chosen and played according to the beliefs on the current trial; Mathys et al., 2014). Our response model was the unit-square sigmoid observation model for binary responses (Iglesias et al., 2013; Mathys et al., 2014). This model estimates on each trial *k* the probability that the agent’s response y is either 0 or 1 (**Figure 2B**; p[y^(k)^ = 1] and p[y^(k)^ = 0]), as a function of the predicted probability that the icon/sequence is rewarding. This mapping from beliefs to decisions depends on the response parameter ζ (interpreted as inverse decision noise). Higher ζ values indicate a greater probability for the agents to select the option that is more likely to be rewarding according to their beliefs. Simulations demonstrate that ζ is recovered well (Hein et al., 2021).

In the following, as stimuli (red and blue icons) are one-to-one associated with motor sequences (seq1 and seq2, respectively), we will use the term action-reward contingency when referring to stimulus-reward or stimulus-outcome mappings.

### Models and priors

In line with previous work (Iglesias et al., 2013; Hein et al., 2021) we fitted the empirical data with different models. We started by modelling our data with the HGF_3_ perceptual model + sigmoid response model, as described above. In this model, the third hierarchical level represents environmental volatility, that is the rate of change in the action-reward contingencies. In our paradigm the true volatility was constant across participants, as the reward contingencies changed approximately every 25-35 trials. The HGF_3_ led to numerical instabilities in 20% of our participants across all groups, in particular in those exhibiting high win rates and thus learning well. The numerical instabilities manifested both when using tight priors (small variance in the prior distribution of *ω*_2_, *ω*__3__) and loose priors (as default in the HGF toolbox); and when using prior values defined in previous work (Iglesias et al., 2013; Hein et al., 2021) or estimated in our data using an ideal observer model. An ideal observer is typically defined as the set of parameter values that minimise the overall surprise that an agent encounters when processing the series of inputs (see an application of an ideal observer model in e.g., Weber et al., 2020). We therefore proceeded to use the 2-level HGF (HGF_2_), in which beliefs on volatility on the third level are fixed. Priors for the perceptual HGF_2_ model were chosen by simulating an ideal observer receiving the series of inputs that the participants observed (**Table 3**). We then used the estimated posterior values on those model parameters as priors for the HGF_2_ perceptual model coupled with our response model. Complementing the HGF, we used two standard reinforcement learning models, the Rescorla-Wagner model (RW; fixed learning rate determined by PEs; Rescorla & Wagner, 1971) and Sutton K1 model (SK1; flexible learning rate driven by recent PEs; Sutton, 1992). Priors for reinforcement learning models were set according to previous literature (Diaconescu, 2014; Hein et al., 2021).

The different models (HGF_2_, RW, SK1) were fitted to the trial-by-trial inputs and responses in each participant using the HGF toolbox, which generates maximum-a-posteriori (MAP) parameter estimates in each individual. To identify the model that explained the behavioural data across all participants best, we used random effects Bayesian model selection (BMS, through the freely available MACS toolbox https://github.com/JoramSoch/MACS; Soch & Allefeld, 2018). Importantly, we used the same priors in all participant groups (HYA, HOA, PD) as in previous studies (Powers et al., 2017; Hein et al., 2021). Note, however, that recent computational modelling work suggests that using different prior values in each participant group may be more suitable to capture between-group differences (e.g., for mental health: Valton et al., 2020).

### Behavioural and computational data analysis

First, we validated the task by assessing the percentage of trials that each sequence type was played (percPlayed). If participants made their decisions based on the inferred contingency mappings, then on average they should have played each sequence type in 50% of the trials. We additionally examined the percentage of trials in which each sequence type was played without performance execution errors (percCorrectlyPlayed).

General task performance in each participant was assessed by analysing the average of the trial-by-trial performance tempo (mIKI in ms: trial-wise mean of the three inter-keystroke-intervals [IKI] across four key presses within the same trial; **Figure 1D**), the mean of the trial-by-trial reaction times (RT in ms: trial-wise time interval between fractal presentation and first key press), percentage of errors (percError: rate of sequences with performance execution errors due to one or several wrong key presses) and win rate (percWin: rate of trials in which the rewarded sequence is played without execution errors). Finally, we also assessed the number of sequence renditions that participants completed during the familiarisation phase (rendFam: average of renditions across both sequence types). Trials with performance execution errors were excluded from analyses on performance tempo to avoid potential confounds, such as slowing following errors (Herrojo Ruiz et al., 2009).

Next, to investigate decision-making processes we analysed group effects on three computational variables that characterised learning in each individual. The model that best explained the behavioural data across all participants according to BMS was the HGF_2_ (see Results section). We therefore assessed the perceptual model parameter *ω_2_* (subject-specific tonic volatility, which influences the speed of belief updating on level 2), ζ (the decision noise of the response model), and the average across trials of *σ*_2_ (posterior variance of the belief distribution). The quantity *σ*_2_ is particularly interesting, as it represents informational uncertainty about the tendency of the action-reward contingency. Moreover, beliefs on level 2 are updated as a function of PEs about the stimulus-outcome mapping (the mismatch between the observed outcomes *u* = 1 or 0 and the agent’s beliefs about the probability of such an outcome) and weighted by *σ*_2_ (the precision ratio on level 2). Accordingly, if agents are more uncertain about the contingencies governing their environment, they will rely more on PEs to update their beliefs on that level.

Finally, to test our main research hypothesis that the strength of expectations about the action-reward contingency modulates the trial-by-trial motor performance, as a function of the group, we focused on the trajectory *µ*_2_ (dropping trial index *k* for simplicity; predictions about the tendency of the action-reward contingency).

### Statistical analyses

#### Bayesian analysis of general task performance and computational variables

First, we calculated the mean and SEM as summary statistics for each of our general task performance (mIKI, RT, percError, percWin, rendFam) and computation variables (*ω*__2__, ζ, *σ*__2__). Next, we evaluated between-group differences by computing Bayes Factors (BF) using the bayesFactor toolbox (https://github.com/klabhub/bayesFactor) in MATLAB. This toolbox implements tests that are based on multivariate generalisations of Cauchy priors on standardised effects (Rouder et al., 2012). In the main study, for each dependent variable (DV), we calculated the BF on two models: m1) DV ∼ 1 + group, dependent variable explained by a fixed effect of group (HYA, HOA, PD); m2) DV ∼ 1 + group + (1|subject), which added the random effect of subjects. Models were fitted using the fitlme function of the MATLAB Statistics toolbox. Computing BF allowed us to quantify the evidence in support of the alternative hypothesis (full models m1 and m2, in our case assessing the main effect of the group) relative to the null model (intercept-only model, i.e., DV ∼ 1). BF values were interpreted as in Andraszewicz et al. (2015). As BF is the ratio between the probability of the data being observed under the alternative hypothesis and the probability of the same data under the null hypothesis, a BF of 20 would indicate strong evidence for the alternative hypothesis. On the other hand, BF of 0.05 would provide strong evidence for the null hypothesis (see Table 1 by Andraszewicz et al., 2015 for further details). For each response variable, m1 was more informative (i.e., provided more evidence either for the null or alternative hypothesis) than m2, and it was also the most parsimonious model among both. Thus, in the following we only reported BF values on m1. Accompanying the BF results, we provided the outcomes of standard one-way analysis of variance (ANOVA) for completion. In the case of main effects being observed in the group-level BF analysis, we conducted follow-up BF analyses on independent two-sample t-tests.

When analysing RT, we first detected and excluded outliers (RT values larger than three standard deviations above the mean) at the subject level. For BF analyses, we used the individual average across 180 trials for the mIKI, RT and *σ*_2_ variables. As mIKI and RT were not normally distributed, values were log-transformed (natural logarithm, log_mIKI, log_RT). The number of renditions during the familiarisation phase was averaged between both types of sequence.

Sanity checks were performed to assess that participants chose to play each sequence as a function of the inferred action-reward contingencies and not based on individual sequence preferences. These were carried out by computing mean and SEM along with BF analyses for paired t-tests on the percentage of trials each sequence type was (correctly) played (percPlayed; percCorrectlyPlayed; outcomes of standard paired t-test reported for completion).

#### Assessing the association between predictions about the action-reward contingency and motor performance using Bayesian Linear Mixed Models

Our main goal was to investigate whether trial-by-trial sequence performance tempo (mIKI) is modulated by the expectation about the tendency of the action-reward contingency (*µ*_2_) in our participant groups. In addition, we aimed to determine whether the group factor modulated the sensitivity of performance tempo to *µ*_2_, resulting in different slopes of the association.

We addressed these questions by implementing a series of Bayesian Linear Mixed Models (BLMM) in R (version 4.0.3). We used the Bayesian Regression Models using Stan (brms; Bürkner, 2017; 2018; 2021) package, freely available on https://cran.r-project.org/web/packages/brms/index.html. Brms relies on the probabilistic programming language Stan, which implements Bayesian inference using Markov Chain Monte Carlo (MCMC) sampling methods to estimate approximate posterior probability distributions for model parameters.

In the HGF for binary categorical inputs, the sign of *µ̂*_2_ (and similarly *µ*_2_) is not informative, as it represents the tendency of an action-reward mapping for an *arbitrary* action (e.g., for seq1). Yet, we could similarly define the model in reference to the other action (e.g., seq2). In line with previous work (Stefanics et al., 2018; Hein et al., 2022), we therefore took the absolute value of *µ̂*_2_ (|*µ̂*_2_|) for our analysis to represent the strength of predictions about the tendency of the action-reward mapping. Trials with greater |*µ̂*_2_| values are trials in which the participants will have a stronger expectation of receiving a reward, given they select the correct action. Thus, |*µ̂*_2_| represents the *strength* of the predictions. We excluded |*µ̂*_2_| values larger than four standard deviations above the mean (0.0017% values removed). Next, we transformed the |*µ̂*_2_| values to z-scores (|*µ̂*_2_|_z) to get a reasonable estimate on the intercept along the mIKI axis. Using |*µ̂*_2_|_z guarantees that the intercept estimate for mIKI reflects the average |*µ̂*_2_| value. As for Bayesian ANOVAs (see Bayesian analysis of general task performance and computational variables), mIKI was log-transformed to approach normality (log_mIKI). In one HOA participant, two log_mIKI values were discarded from analyses as they were not registered correctly in the JSON file (i.e., represented an impossible value of mIKI ∼ 50 ms). In BLMM with brms, it is standard to select one group as reference for the parameter estimates. Brms then estimates the posterior distribution of parameter differences between each group and the reference group, as well as the posterior distributions of parameters in the reference group itself. We set HOA as the reference group, and therefore posterior distributions of between-group differences on response variables were assessed for HOA vs HYA and HOA vs PD.

We implemented six models of increasing complexity, with every model including a larger number of explanatory variables (**Table 4**). For simplicity, in the following we used variable label y to represent our dependent variable log_mIKI, and x to represent the explanatory variable |*µ̂*_2_|_z. To answer our research questions, we primarily focused on: (i) the fixed effect of x (sensitivity [slope] of the performance tempo to the strength of predictions about the action-reward contingency in the reference group, HOA); and (ii) the interaction effect x * group (differences between groups in the sensitivity [slope] of the performance tempo to the strength of expectations about the action-reward mapping).

**Table 4.**
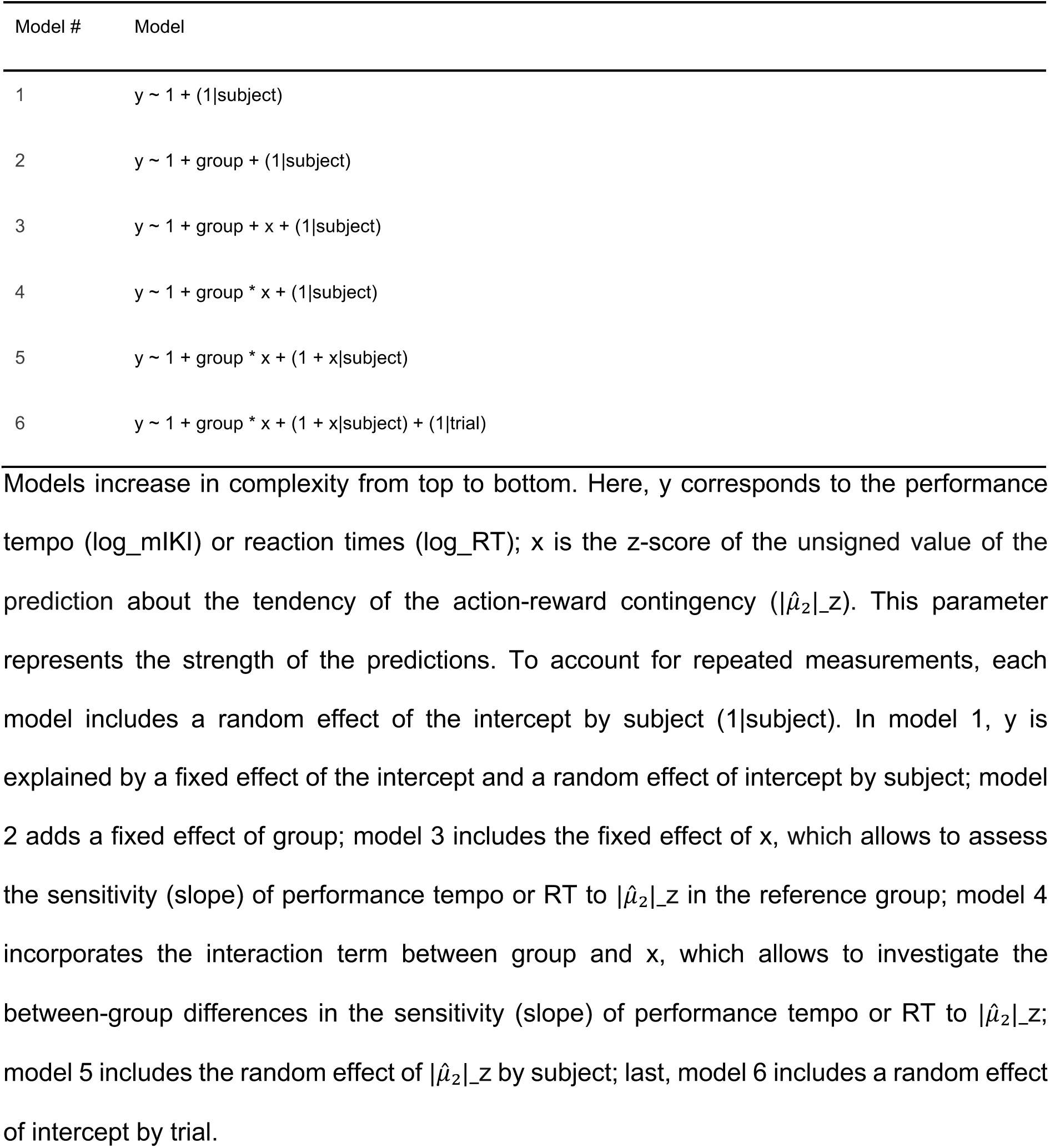
Models of increasing complexity used to explain the modulation of motor performance by the strength of the expectation about the tendency of the action-reward contingency.

For each model we ran four independent chains with 5000 iterations each, of which the first 1000 were discarded as warmup. This resulted in a total of 16000 posterior samples. In all models we used default prior distribution for the intercept, and a normal distribution for each fixed and random effect (fixed effects for group and x, normal [0,2)]; interaction term group * x, normal [0,1]; random effects for intercept by subject and intercept by trial, normal [0,2]; random effect x by subject, normal [0,1]). The prior on the LKJ-Correlation, the correlation matrices in brms (Lewandowski, Kurowicka, & Joe, 2009), was set to 2 as recommended in Bürkner and colleagues (2017). Chain convergence was assessed using the Gelman-Rubin statistics (R-hat < 1.1; Gelman and Rubin, 1992).

Models were compared using leave-one-out cross-validation of the posterior log-likelihood (LOO-CV) with Pareto-smoothed importance sampling (Vehtari et al., 2017). The identification of the best fitting model was based on the highest expected log point-wise predictive density (ELPD). We also checked that the absolute mean difference in ELPD between two models (elpd_diff in brms) exceeded twice the standard error of the differences (2 * se_diff). LOO-CV identified the most complex model (model number 6 in **Table 4**) as the best fitting model (see Results section for further details). This model explained the performance tempo as the interaction between group and the strength of the expectation about the action-reward contingency (in addition to main effects). Further, it modelled the effect of subjects on the intercept and |*µ̂*_2_|_z as a random effect, and the effect of trials on the intercept as a random effect. In the results section, we reported for each parameter the posterior point estimate and the associated 95% credible interval (CI).

Because reward expectations could modulate RT as shown previously (Codol et al., 2020), we conducted an additional analysis to assess the effect of |*µ̂*_2_| on RT trial-by-trial. Further, we evaluated whether the group factor influences the sensitivity of RT to |*µ̂*_2_|. In this analysis, we followed the same procedure as for the sequence performance tempo analysis. In particular, the association between RT (log-transformed) and |*µ̂*_2_|_z was assessed by implementing and comparing six models of increasing complexity in brms (**Table 4**; see results for further details). RT values smaller than 200 ms, larger than 2500 ms or three standard deviations above the mean were excluded from statistical analyses. As for performance tempo, in the results section we used the variable label y for the dependent variable log_RT and x for |*µ̂*_2_|_z.

#### Bayesian analyses on the control study

As described above, in the control study participants were allocated to two different analysis groups (Q8_T_ and Q8_F_) depending on their answer to a post-performance question (“I could *always* distinguish whether 0 points reflected a performance error or a bad decision”, binary answer: True/False). This allowed us to test the potential role of subjective inferences about task-related reward assignment in explaining our main results.

As for the main experiment, we computed the mean and SEM as summary statistics for each dependent variable. Next, we used the bayesFactor toolbox to calculate the evidence in support of (or against) group differences in general task performance (mIKI, RT, percError, percWin) and computational variables (*ω*__2__, ζ, *σ*__2__). We intentionally did not analyse the rate of sequence renditions during the familiarisation phase as here we were only interested in assessing the role of subjective inferences about credit assignment on motor sequence performance decision-making behaviour. We performed BF analysis on independent two-sample t-tests to assess between group-differences on the variables of interest (results on standard independent t-tests also reported for completion). As described above, RT values larger than three standard deviations above the mean were discarded. Prior to BF analysis, the values of mIKI, RT and *σ*__2__ variables were averaged across the 180 trials within each participant. Also, mIKI and RT were log-transformed (log_mIKI, log_RT) to approach normality.

Next, to test potential between-group differences in the mIKI-|*µ̂*_2_| association, we implemented six BLMM of increasing complexity (same models as in the main experiment, **Table 4**). We excluded |*µ̂*_2_| values larger than four standard deviations above the mean (0.0006% values removed) and mIKI was log-transformed (log_mIKI). Moreover, the unsigned |*µ̂*_2_| values were transformed into |*µ̂*_2_|_z. As for the main experiment, the most complex model (model number 6 in **Table 4**) was identified as the best fit by LOO-CV (see Results section for further details). As the results in the main experiment showed no modulation of RT by the strength of predictions about the action-reward contingencies, we did not investigate a potential RT-|*µ̂*_2_| association in the control sample.

Finally, we evaluated whether Q8_T_ and Q8_F_ differed in the rate of subjective number estimate of performance errors. In particular, we were interested in assessing between-group differences in the tendency of under/overestimating the number of performance errors. For each participant, the rate of subjective performance execution errors (subjective_percError) was calculated through the post-performance questionnaire (see Questions 1,2,3 **Table 2**). We arbitrarily assigned a value of 0.028 (= 5/180) if subjects thought to have committed less than 10 performance errors; 0.111 (= 20/180) for between 20 and 40 estimated performance errors; 0.222 (= 40/180) for more than 40 subjective performance errors. To assess whether this rough estimate of the percentage of performance errors reflected a general over or underestimation of the true performance error rate in the total sample (N = 39), we first conducted a BF analysis on the correlation between the subjective and empirical error rates (Pearson’s r coefficient and p-value reported for completion). Next, we identified potential group-related systematic biases in the subjective estimate. This was done with a BF analysis using independent two-sample t-tests on the normalised rate of subjective errors ([subjective_percError-percError]/percError; results on standard independent t-tests reported for completion).

## RESULTS

### General task performance

Before assessing between-group differences on the main markers of general task performance (mIKI, RT, percError, percWin, rendFam), we validated our task by calculating the rate of performance for each sequence type. Participants played on average seq1 and seq2 50% of the trials (seq1: mean 0.490, SEM 0.008; seq2: mean 0.508, SEM 0.008). Indeed, despite participants committing fewer performance execution errors for seq1 (mean 0.958, SEM 0.005) than seq2 (mean 0.922, SEM 0.008; percCorrectlyPlayed, BF = 1126.7, suggesting extreme evidence for alternative hypothesis that the rate of correct performance differed in seq1 and seq2, t_(93)_ = 4.576, p < 0.001), they did not express a preference towards a sequence type (percPlayed, BF = 0.2295, moderate evidence in support of the null hypothesis for no differences in the percentage of performances by sequence type, t_(93)_ = - 1.204, p = 0.232).

Overall, as expected, our analyses revealed between-group differences in average performance tempo (mIKI in ms, HYA: 300, SEM:15.8; HOA: mean 424, SEM 19.6; PD: mean 537, SEM 26.9; **Figure 3A**) and reaction time (RT in ms, HYA: 669, SEM: 35.5; HOA: mean 888, SEM 49.5; PD: mean 979, SEM 77.2; **Figure 3B**), with movements progressively slowing down in ageing and PD patients. BF analyses on mean performance tempo yielded extreme evidence for a group effect (log_mIKI: BF = 1.1253e+09, demonstrating extreme evidence for the alternative hypothesis; F_(2,91)_ = 35.332, p < 0.001). Post hoc pair-wise t-tests using BF showed extreme evidence for between-group differences in HYA vs HOA (BF = 1.2044e+04) and in HYA vs PD (BF = 3.3592e+07). We also found very strong evidence for the alternative hypothesis in HOA vs PD (BF = 32.591). Thus, average performance tempo was differently modulated between groups, with HYA being faster than HOA and PD, and HOA faster than PD. Regarding RT, there was extreme evidence supporting between-group differences (log_RT: BF = 282.230; F_(2,91)_ = 10.893, p < 0.001). BF analysis on post hoc independent two-sample t-tests revealed very strong and extreme evidence for between-group differences in HYA vs HOA (BF = 80.744) and HYA vs PD (BF = 183.289), respectively. Yet, we only found anecdotal evidence in support of the null hypothesis in HOA vs PD (BF = 0.404). Hence, despite HYA displaying shorter RTs than HOA and PD, our analyses suggest similar RTs in HOA and PD.

**Figure 3.**
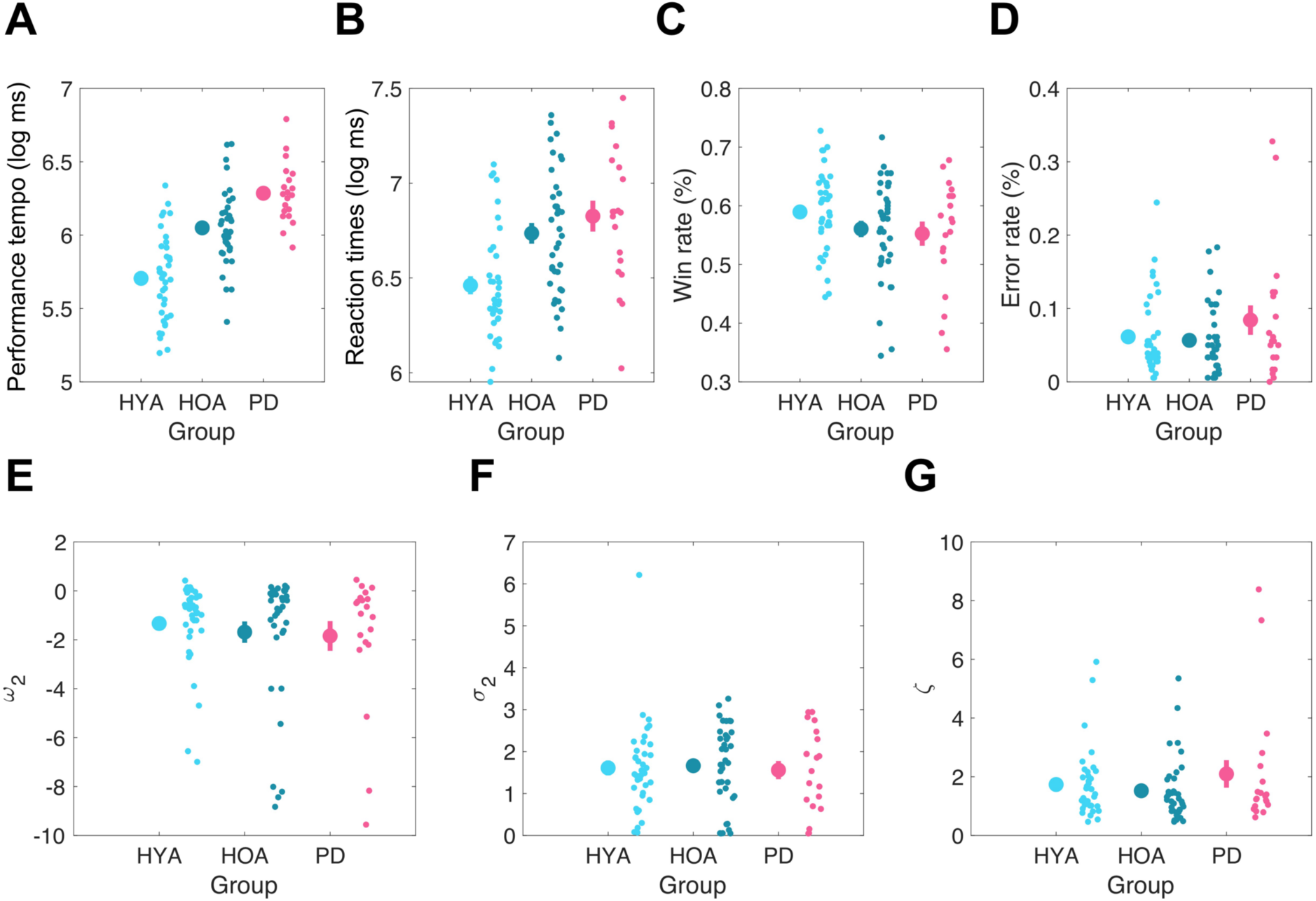
Markers of general task performance and decision making in healthy younger adults (HYA; in light blue), healthy older adults (HOA; in dark blue) and patients with Parkinson’s Disease (PD; in purple). ***A***, Performance tempo (mIKI, mean inter-keystroke-interval, in ms); ***B***, Reaction time (RT, in ms); ***C***, Rate of win trials (percWin); ***D***, Rate of performance execution errors (percError); ***E***, tonic volatility (*ω*_2_); ***F***, informational uncertainty on level 2 (*σ*_2_); ***G***, response model parameter (ζ). Values mIKI, RT and *σ*_2_ are averaged across 180 trials within each participant. mIKI and RT values are log-transformed. In every plot, to the right of each mean (large dot) and standard error of the mean (denoted by the vertical bar) the individual data points in each group are shown to visualise group population variability.

In addition, we found anecdotal evidence supporting that groups differed in the number of sequence renditions during the familiarisation phase (rendFam, HYA: mean 5.6, SEM 0.1; HOA: mean 6.0, SEM 0.2; PD: mean 7.1, SEM 0.8; BF = 1.733; F_(2,91)_ = 4.448, p = 0.014).

Post-hoc BF analyses to assess differences between pairs of groups revealed anecdotal and moderate evidence for between-group differences in HYA and HOA (BF = 1.900) and HYA and PD (BF = 3.030), respectively. Still, HOA and PD practised the two sequences to a similar extent (BF = 0.853, revealing anecdotal evidence for the null hypothesis). Of note, practising more during familiarisation was not associated with better win rates or average performance tempo during task completion. A correlation analysis across all participants between the number of repetitions during familiarisation and these variables demonstrated some evidence for null correlation effects (percWin: BF = 0.290, Pearson r = -0.134, p = 0.200; log_mIKI: BF = 0.397; Pearson r = 0.158, p = 0.131; note that we excluded one PD patient who practised 21 times during familiarisation as outlier in this correlation analysis).

The group effects observed above were not accompanied by a dissociation between groups in the win rate or the rate of performance execution errors (**Figure 3C-D**). BF analysis on win rates provided moderate evidence for the lack of a group effect (percWin, HYA: mean 0.590, SEM 0.012; HOA: mean 0.561, SEM 0.014; PD: mean 0.553, SEM 0.021; BF = 0.210, supporting moderate evidence for the null hypothesis; F_(2,91)_= 1.848, p = 0.163). A similar outcome was observed in the analysis of performance execution error rates (percError, HYA: mean 0.061, SEM 0.009; HOA: mean 0.057, SEM 0.008; PD: mean 0.084, SEM 0.020; BF = 0.146, moderate evidence for the null hypothesis; F_(2,91)_ = 1.456, p = 0.239). In sum, we found moderate evidence that HYA, HOA and PD did not differ in neither the rate of win nor error trials.

### Computational parameters

Decision making was assessed by looking at between-group differences in the computational variables *ω*_2_, ζ and *σ*_2_. After excluding the HGF_3_ from model comparison due to numerical instabilities, BMS was conducted on the HGF_2_ and two reinforcement learning models (RW, SK1) using the individual log-model evidence (LME) values provided by the HGF toolbox. The winning model was the HGF_2_, with an exceedance probability of 0.95 and an expected frequency of 0.90. Of note, although the HGF_3_ model was not included in BMS, a qualitative comparison of LME values for the HGF_3_ and HGF_2_ models in the 80% participants in which HGF_3_ did not lead to numerical instabilities revealed extremely similar values (LME differences < 1). This observation suggested that both models described behaviour in our task with constant true volatility to a similar degree.

Overall, we found no group effect on the signatures of reward-based learning and decision making in our volatile task (**Figure 3E-G**). BF analysis on ’_2_ demonstrated strong evidence for the absence of a main effect of group (HYA: mean -1.332, SEM 0.282; HOA: mean -1.686, SEM 0.438; PD: mean -1.843, SEM 0.609; BF = 0.059; F_(2,91)_ = 0.380 p = 0.685). Similarly, we found strong evidence in favour of a lack of group effect on the informational uncertainty about beliefs on the tendency of the action-reward contingency, *σ*_2_ (HYA: mean 1.610, SEM 0.177; HOA: mean 1.663, SEM 0.158; PD: mean 1.559, SEM 0.218; BF = 0.045; F_(2,91)_ = 0.074, p = 0.928). Last, groups exhibited a similar mapping from beliefs to responses, driven by the response model parameter ζ (HYA: mean 1.735, SEM 0.191; HOA: mean 1.523, SEM 0.176; PD: mean 2.095, SEM 0.469; BF = 0.114, demonstrating moderate evidence for the null hypothesis; F_(2,91)_ = 1.1495, p = 0.321).

A direct comparison between the Italian HOA subsample and (Italian) PD sample revealed anecdotal or moderate evidence in support of the null hypothesis when assessing general performance and decision-making variables (exception for log_mIKI). These findings thus converge with the outcomes of the full HOA sample analysis. On the other hand, the very strong evidence in support of group effects on the performance tempo in the full sample was only anecdotal when directly comparing Italian HOA and PD samples on this variable (log_mIKI: BF = 2.556; t_(42)_ = -2.348, p = 0.024). These results suggested that Italian healthy ageing was associated with slower performance tempo relative to UK healthy ageing participants (log_mIKI: BF = 6.637; t_(35)_ = 2.871, p = 0.007; moderate evidence supporting differences in performance tempo).

### Sensitivity of motor performance to the strength of expectations about the action-reward contingency

For performance tempo, LOO-CV identified the most complex model (model number 6) as the best fit (**Table 5**). Posterior predictive checks revealed that the best model had strong predictive power for the range of the DV (**Figure 4A**). In the following we use variable label y to represent our dependent variable log_mIKI (in log-ms), and x to represent the explanatory variable |*µ̂*_2_|_z.

**Figure 4.**
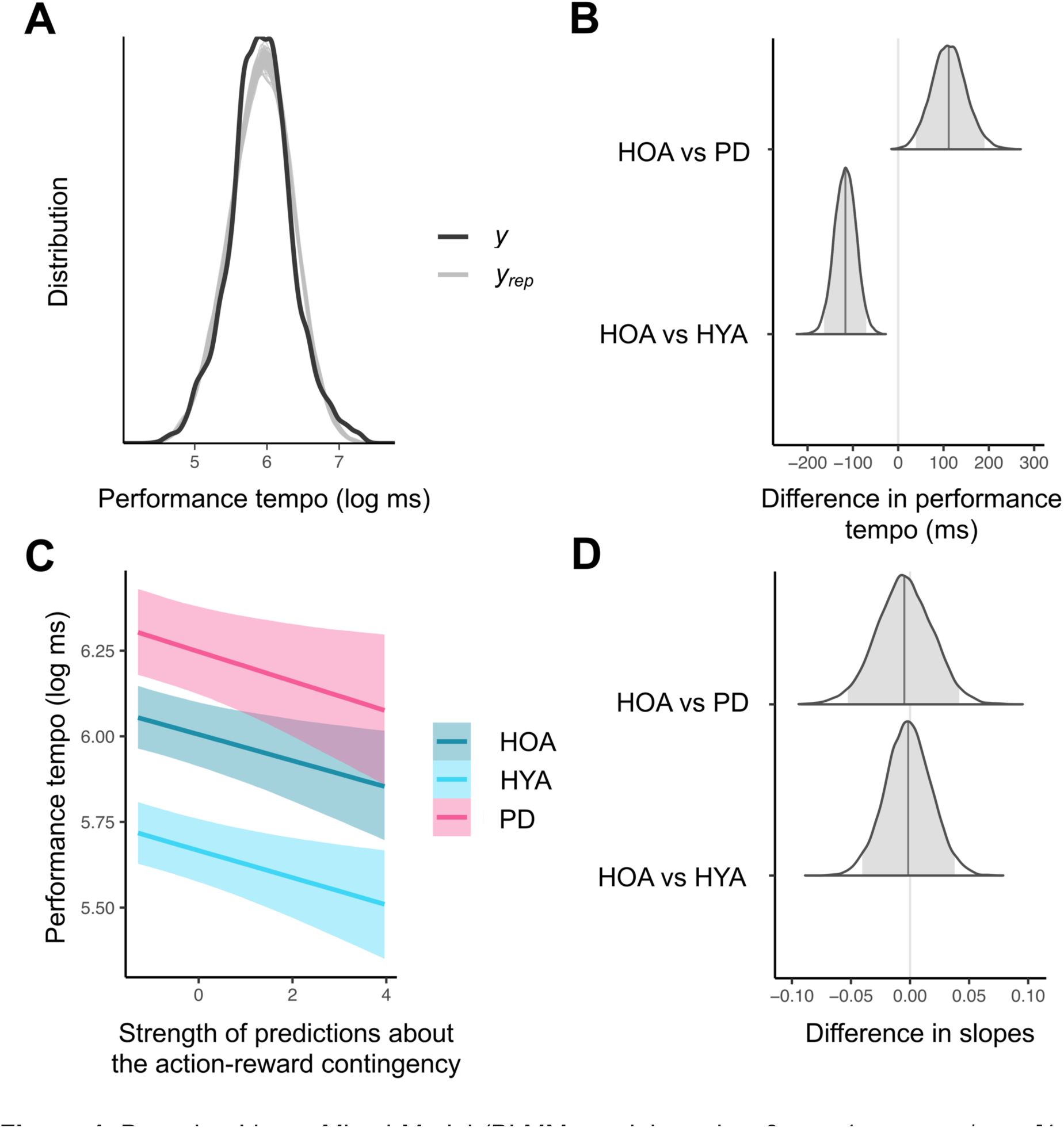
Bayesian Linear Mixed Model (BLMM; model number 6, y ∼ 1 + group * x + [1 + x|subject] + [1|trial]) with healthy older adults (HOA) as the reference group. ***A***, Illustration of the posterior predictive checks where the distribution of the observed outcome variable (y, in our case performance tempo) is compared to simulated datasets (y_rep_) from the posterior predictive distribution (100 draws). ***B***, Distributions of the difference in ms between performance tempo (intercept) in HOA and healthy younger adults (HYA), and in HOA and patients with Parkinson’s Disease (PD). For each distribution, the grey vertical bar indicates the posterior point estimate, while the grey area under the curve represents the 95% credible interval (CI). In the current plot, CIs do not overlap with zero (the null hypothesis). This indicates that there is a 95% probability of between-group differences in performance tempo. ***C***, Results of the BLMM analysis. We analysed how the strength of predictions about the action-reward contingency modulates performance tempo separately for HYA (in light blue), HOA (in dark blue) and PD (in purple). Here, mIKI (performance tempo: mean inter-keystroke- interval) values are represented in the log-scale. The negative slopes suggest that stronger predictions about the action-reward contingency are associated with faster performance tempo. ***D***, Distributions of the difference between slopes in HOA vs HYA, and HOA vs PD. Here, as CIs include zero we can conclude with 95% probability that groups do not differ in how the strength of predictions about the reward contingency influences motor performance tempo. Thus, the sensitivity of performance tempo to the strength of predictions about the reward mapping is not differently modulated between groups.

**Table 5.**
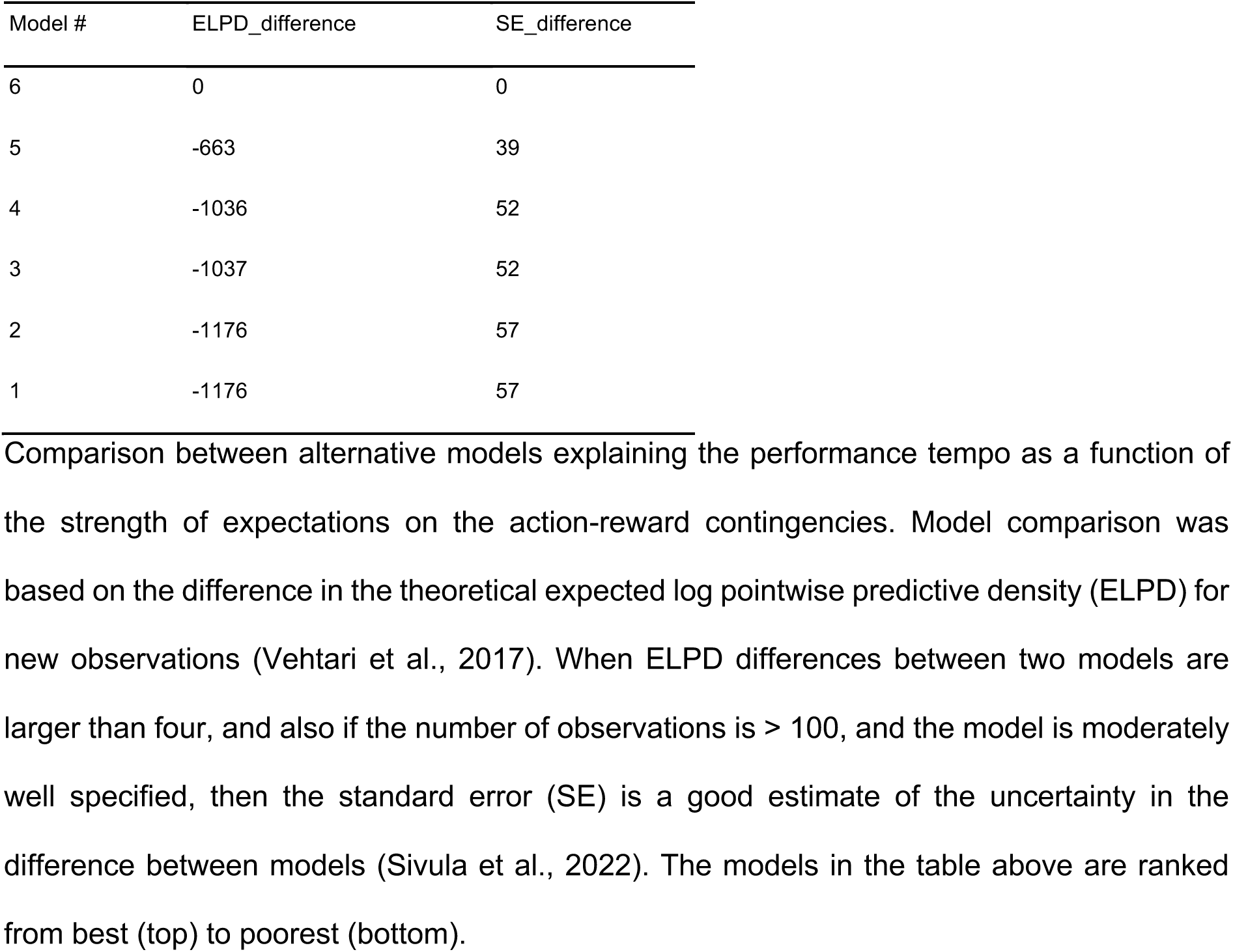
Assessment of predictive accuracy in the models on performance tempo.

First, we found that groups differed in performance tempo, as expected. This is in line with our previous between-group analyses showing a progressive slowness in execution tempo in HOA and PD. The posterior estimate for the intercept in the reference group, HOA, was 6.01, CI = [5.91, 6.10] (in ms, 406, CI = [369, 445]). This corresponds to the posterior distribution of y when x values are equal to zero. The distribution of the differences between intercepts in HOA and HYA had a posterior estimated value of -0.34, CI = [-0.47, -0.21] (in ms, -117, CI = [-164, -70]), while the distribution of the differences between intercepts in HOA and PD yielded a posterior point estimate of 0.24, CI = [0.09, 0.40] (in ms, 112, CI = [39, 191]). As neither of the two distributions overlapped with zero, we concluded that HYA performed the sequences faster than HOA (HOA vs HYA distribution is shifted towards the negative x-axis), while PD was slower than HOA (HOA vs PD distribution is shifted towards the positive x-axis) (**Figure 4B**).

Next, we evaluated how the strength of predictions about the action-reward contingency modulated performance tempo on a trial-by-trial basis. The analyses supported our hypothesis, showing that stronger expectations about the reward contingency invigorated motor performance through faster execution tempo. Here, we focused on the distribution of the fixed effect of x (slope of the association between y and x) in the reference group, HOA. This distribution informs about the sensitivity of the performance tempo to the strength of predictions about the action-reward contingency in HOA. The posterior estimate of x was equal to -0.04, CI = [-0.07, -0.01]. As the distribution did not include zero, this highlights a negative relationship between performance tempo and the strength of expectations about the action-reward contingency in the reference group (**Figure 4C**).

As we were also interested in evaluating potential between-group differences in the association (slope) between the strength of predictions about the action-reward contingency and performance tempo, we assessed the distribution of the interaction effect group * x. Both the posterior distributions of slope differences between HOA and HYA and between HOA and PD overlapped with zero, suggesting that the sensitivity of performance tempo to the strength of expectations about the action-reward contingency did not differ between groups (HOA vs HYA: posterior estimate = 0, CI = [-0.04, 0.04]; HOA vs PD: posterior estimate = 0, CI = [-0.05, 0.04]; **Figure 4D**).

Finally, the estimate of the standard deviation for the random effect of intercept by subject was 0.28, CI = [0.25, 0.33], providing a measure for inter-individual intercepts variability in performance tempo. The estimate of the standard deviation for the random effect of x by subject amounted to 0.08, CI = [0.07, 0.10] and informed about inter-individual differences in the sensitivity of performance tempo to the strength of predictions about the reward contingency. Finally, the estimate of the standard deviation for the random effect of intercept by trial was 0.08, CI = [0.07, 0.09], signalling trial-by-trial differences in performance tempo. Participants indeed played the sequences faster towards the end of the experiment, reflecting a practice effect (**Figure 1D**).

Overall, these results (see **Table 6** for a summary of the posterior distributions for the winning model) demonstrated that motor performance tempo was influenced trial-by-trial by the strength of predictions about the tendency of the action-reward contingency, with stronger expectations leading to faster execution tempo. Importantly, the sensitivity of performance tempo to the strength of these predictions was not differently modulated between groups, suggesting that all groups could successfully use to a similar degree the inferred predictions to invigorate their motor performance.

**Table 6.**
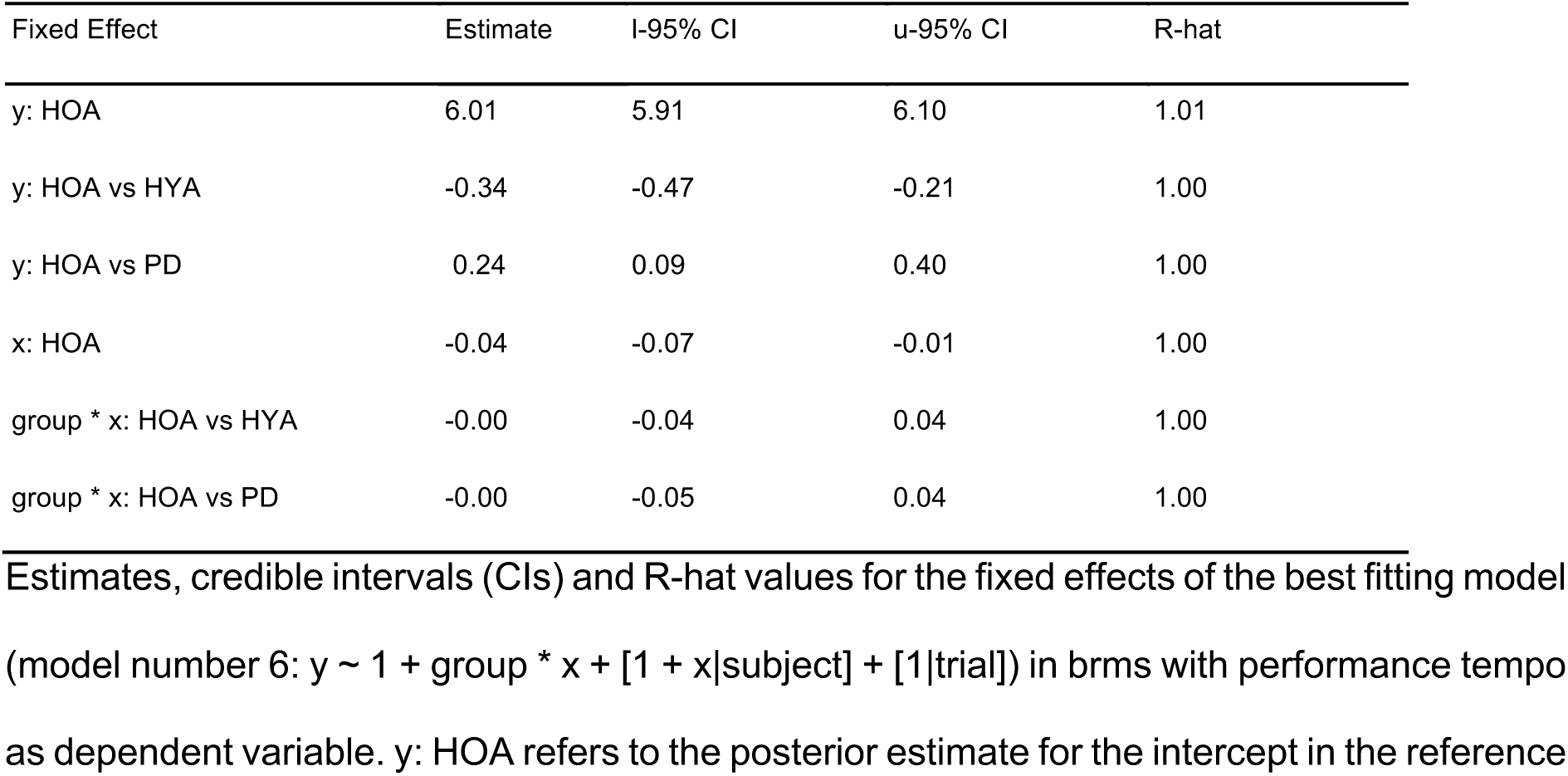
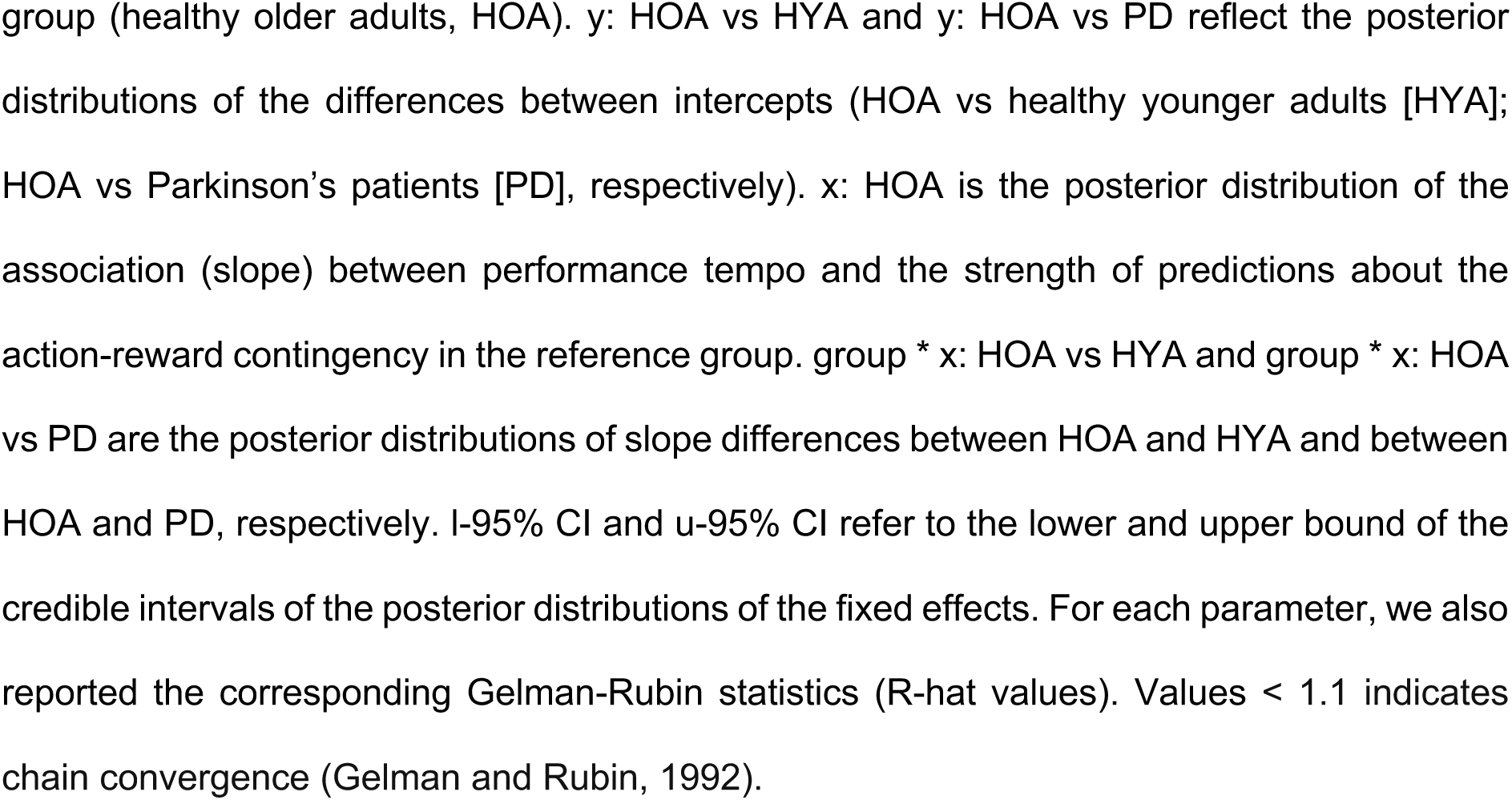
Summary of the posterior distributions for the fixed effects of model number 6 on performance tempo.

We additionally assessed whether the motor invigoration effect extended to the RT, reflecting the time to initiate the sequence performance (first key press). As for performance tempo, LOO-CV identified model number 6 as the best fit (**Table 7**) and posterior predictive checks demonstrated good predictive power for the range of the DV (**Figure 5A**).

**Figure 5.**
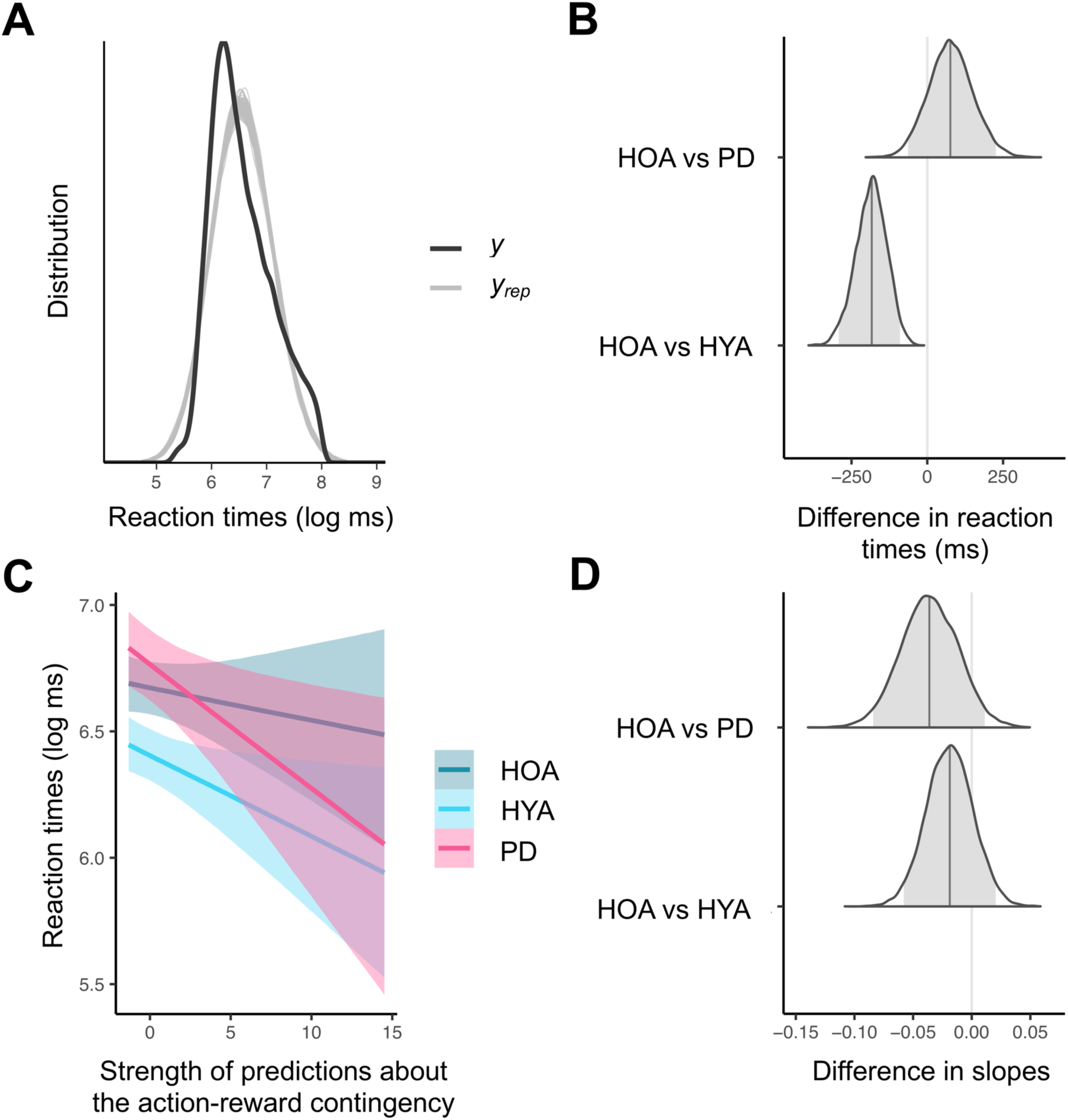
Bayesian Linear Mixed Model (BLMM; model number 6, y ∼ 1 + group * x + [1 + x|subject] + [1|trial]) with healthy older adults (HOA) as the reference group. ***A***, Illustration of the posterior predictive checks where the distribution of the observed outcome variable (y, in our case reaction times [RT]) is compared to simulated datasets (y_rep_) from the posterior predictive distribution (100 draws). ***B***, Distributions of the difference in ms between RT (intercept) in HOA and healthy younger adults (HYA), and in HOA and patients with Parkinson’s Disease (PD). For each distribution, the grey vertical bar indicates the posterior point estimate, while the grey area under the curve represents the 95% credible interval (CI). In the current plot, CI of the bottom distribution does not overlap with zero (the null hypothesis). This indicates that there is 95% probability of between-group differences in RT. On the other hand, the distribution at the top includes zero. This suggests that there is 95% probability of HOA and PD not differing in RT. ***C***, Results of the BLMM analysis. We analysed how the strength of predictions about the action-reward contingency modulates RT separately for HYA (in light blue), HOA (in dark blue) and PD (in purple). Here, RT values are represented in the log-scale. We found no modulation of RT by the strength of expectations about the reward mapping. ***D***, Distributions of the difference between slopes in HOA vs HYA, and HOA vs PD. Here, as CIs include zero we can conclude with 95% probability that groups do not differ in how the strength of predictions about the reward contingency influences RT. Thus, the sensitivity of RT to the strength of predictions about the reward mapping is not differently modulated between groups.

**Table 7.**
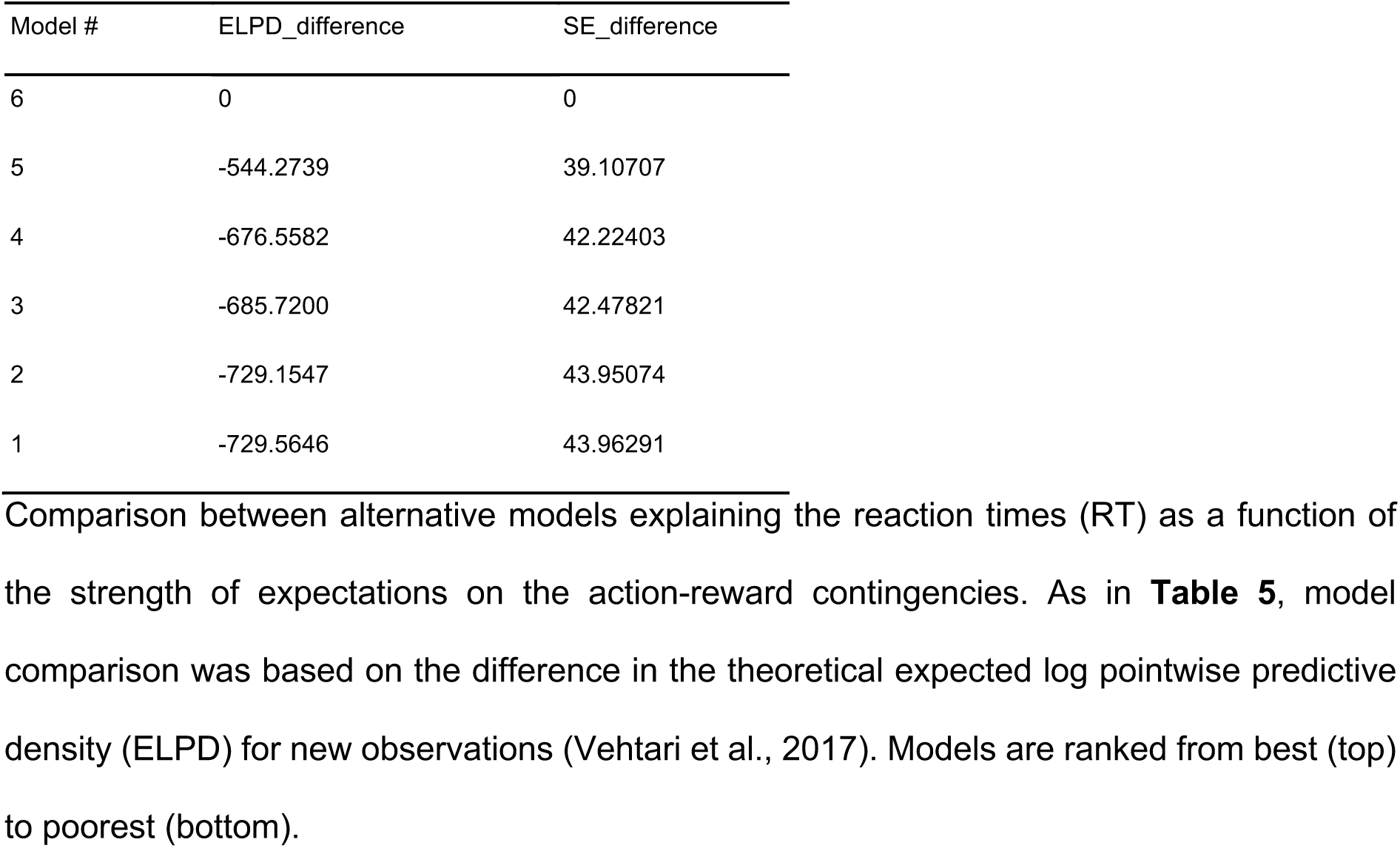
Assessment of predictive accuracy in the models on reaction times.

| Model # | ELPD_difference | SE_difference |
| --- | --- | --- |
| 6 | 0 | 0 |
| 5 | -544.2739 | 39.10707 |
| 4 | -676.5582 | 42.22403 |
| 3 | -685.7200 | 42.47821 |
| 2 | -729.1547 | 43.95074 |
| 1 | -729.5646 | 43.96291 |

Our brms analysis on the best fitting model revealed shorter RT in HYA compared to HOA, with no differences emerging between HOA and PD. The posterior point estimate for the intercept in the reference group, HOA, was 6.67, CI = [6.57, 6.78] (in ms, 791, CI = [712, 876]), which reflects the posterior distribution of y when x values are set to zero. The distribution of the differences between intercepts in HOA and HYA was centred at -0.27, CI = [-0.42, -0.13] (in ms, -185, CI = [-292, -90]), which did not overlap with zero and was shifted towards the negative x-axis (**Figure 5B**). On the other hand, the distribution of the differences between intercepts in HOA and PD yielded a posterior point estimate of 0.09, CI = [-0.08, 0.26] (in ms, 77, CI = [-64, 226]) and included zero. These results demonstrated that HYA had faster RT than HOA, consistent with our mIKI group results, whereas PD and HOA had a similar RT intercept.

Regarding the association between the strength of predictions about the action-reward contingency and RT, we observed no trial-by-trial modulation and no group effects. The distribution of the fixed effect of x (slope of the association between y and x in the reference group, HOA) had a posterior point estimate of -0.01, CI [-0.04, 0.02]. As the distribution’s centre overlapped with zero, this demonstrates that the strength of predictions about the action-reward contingency did not modulate RT in this group (**Figure 5C**). Potential between-group differences in the slope were assessed by investigating the distribution of the interaction effect group * x. Both the posterior distributions of slope differences between HOA and HYA and between HOA and PD included zero (HOA vs HYA: posterior estimate = -0.02, CI = [- 0.06, 0.02]; HOA vs PD: posterior estimate = -0.04, CI = [-0.08, 0.01]; **Figure 5D**). This outcome supported that the sensitivity of RT to the strength of expectations about the reward mapping did not differ between groups. Thus, the strength of predictions about the action-reward contingency invigorated performance tempo on a trial-by-trial basis without affecting the RT (see **Table 8** for a summary of the posterior distributions for the winning model on RT). Also, the estimates of the standard deviation for the random effects of intercept and x by subject were 0.31, CI = [0.27, 0.36] and 0.08, CI = [0.06, 0.09], respectively. The former informed about the inter-individual variability in RT intercepts, while the latter provided a measure of slope differences among individuals between RT and the strength of expectations about the reward mapping. The estimate of the standard deviation for the random effect of the intercept by trial was 0.15, CI = [0.13, 0.17], which reflected the speeding up of RTs across trials.

**Table 8.** Summary of the posterior distributions for the fixed effects of model number 6 on RT.

| Fixed Effect | Estimate | l-95% CI | u-95% CI | R-hat |
| --- | --- | --- | --- | --- |
| y: HOA | 6.67 | 6.57 | 6.78 | 1.01 |
| y: HOA vs HYA | -0.27 | -0.42 | -0.13 | 1.01 |
| y: HOA vs PD | 0.09 | -0.08 | 0.26 | 1.00 |
| x: HOA | -0.01 | -0.04 | 0.02 | 1.00 |
| group * x: HOA vs HYA | -0.02 | -0.06 | 0.02 | 1.00 |
| group * x: HOA vs PD | -0.04 | -0.08 | 0.01 | 1.00 |

Estimates, credible intervals (CIs) and R-hat values for the fixed effects of the best fitting model (model number 6: $y \sim 1 + \text{group} * x + [1 + x|\text{subject}] + [1|\text{trial}]$ ) in brms with reaction times (RT) as dependent variable. y: HOA refers to the posterior estimate for the intercept in the reference group (healthy older adults, HOA). y: HOA vs HYA and y: HOA vs PD reflect the posterior distributions of the differences between intercepts (HOA vs healthy younger adults [HYA]; HOA vs Parkinson's patients [PD], respectively). x: HOA is the posterior distribution of the association (slope) between RT and the strength of predictions about the action-reward contingency in the reference group. group \* x: HOA vs HYA and group \* x: HOA vs PD are the posterior distributions of slope differences between HOA and HYA and between HOA and PD, respectively. l-95% CI and u-95% CI refer to the lower and upper bound of the credible intervals of the posterior distributions for the fixed effects. For each parameter, we also reported the corresponding Gelman-Rubin statistics (R-hat values). Values < 1.1 indicates chain convergence (Gelman and Rubin, 1992).

### Subjective inference about task-related reward assignment

We conducted Bayesian analyses on the HYA control sample to evaluate whether subjective inferences about the hidden causes for the lack of reward could modulate our main results.

Overall, our analyses provided anecdotal and moderate evidence for the lack of differences between Q8_T_ and Q8_F_ in the main markers of general task performance (log_mIKI: BF = 0.417; t_(37)_ = -0.795, p = 0.432; log_RT: BF = 0.331; t_(37)_ = 0.202, p = 0.841; percWin: BF = 0.408; t_(37)_ = 0.758, p = 0.453; percError: BF = 0.596; t_(37)_ = -1.252, p =0.219; see **Figure 6A-D** for summary statistics). Similar results were observed for computational variables, as BF analysis on *ω*_2_, ζ and *σ*_2_ provided anecdotal evidence for the absence of a group effect (*ω*_2_: BF = 0.560; t_(37)_ = -1.183, p = 0.244; ζ: BF = 0.445; t_(37)_ = 0.895, p = 0.377; *σ*_2_: BF = 0.463; t_(,37)_ = -0.951, p = 0.348; see **Figure 6E-G** for summary statistics).

**Figure 6.**
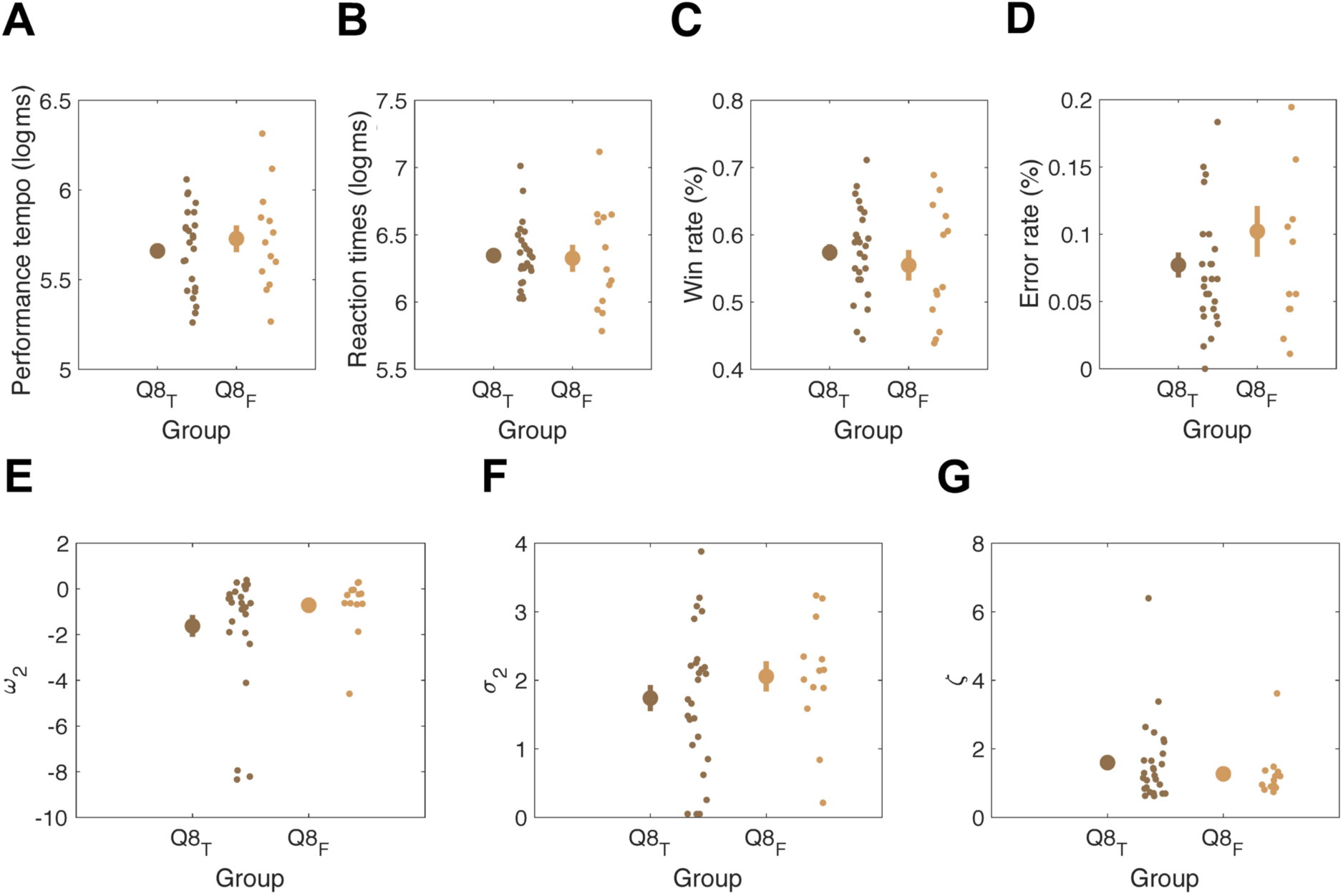
Markers of general task performance and decision making in participants that replied True to Question 8 (Q8_T_; in dark brown) and participants that replied False to Question 8 (Q8_F_; in light brown) in the post-performance questionnaire (see **Table 2**). ***A***, Performance tempo (mIKI, mean inter-keystroke-interval; in ms, Q8_T_: mean 287, SEM 13.2; Q8_F_: mean 307, SEM 27.2); ***B***, Reaction times (RT; in ms, Q8_T_: mean 570, SEM 30.5; Q8_F_: mean 559, SEM 69.7); ***C***, Rate of win trials (percWin; Q8_T_: mean 0.574, SEM 0.013; Q8_F_: mean 0.555, SEM 0.024); ***D***, Rate of performance execution errors (percError; Q8_T_: mean 0.077, SEM 0.010; Q8_F_: mean 0.102, SEM 0.020); ***E***, tonic volatility, (*ω*_2_;Q8_T_: mean -1.624, SEM 0.510; Q8_F_: mean -0.715, SEM 0.357); ***F***, informational uncertainty on level 2 (*σ*_2_; Q8_T_: mean 1.740, SEM 0.203; Q8_F_: mean 2.057, SEM 0.237); ***G***, response model parameter, (ζ; Q8_T_: mean 1.599, SEM 0.237; Q8_F_: mean 1.271, SEM 0.206). Values mIKI, RT and “_2_ are averaged across 180 trials within each participant. mIKI and RT values are log-transformed. In every plot, to the right of each mean (large dot) and standard error of the mean (denoted by the vertical bar) are displayed the individual data points in each group to visualise group population variability.

Hence, whether participants were *always* certain (Q8_T_) or not (Q8_F_) of the implications of receiving zero points, their general motor sequence performance and decision-making behaviour seemed similar, albeit mainly based on anecdotal evidence.

We further investigated whether not being *always* sure about the causes for the lack of reward could impact the sensitivity of performance tempo to the strength of predictions about the action-reward contingency. As for the main experiment, LOO-CV identified the most complex model (model number 6) as the best fit (**Table 9**). Posterior predictive checks demonstrated a very strong predictive power for the range of RT values in the best model (**Figure 7A**).

**Figure 7.**
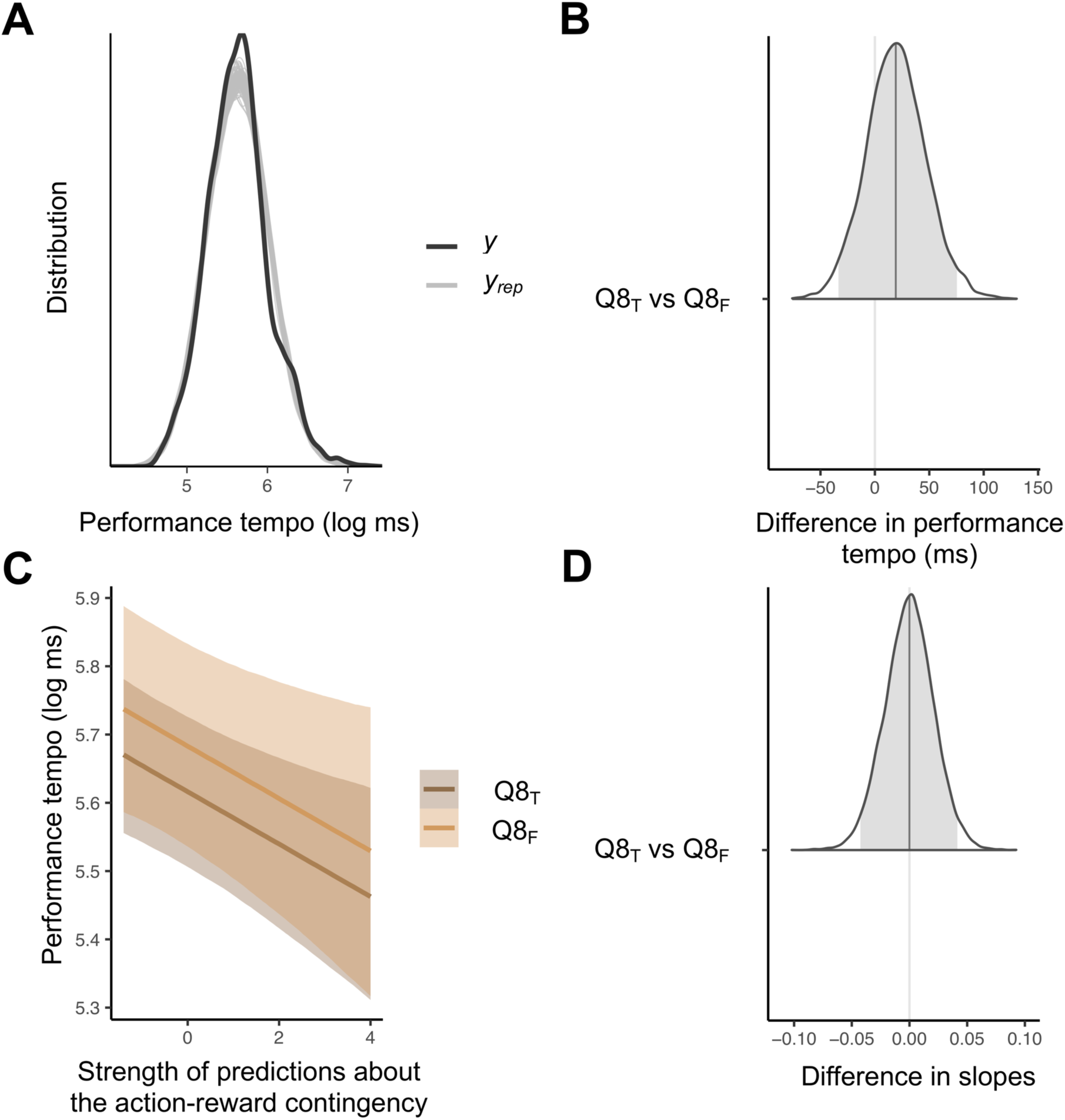
Bayesian Linear Mixed Models (BLMM; model number 6, y ∼ 1 + group * x + [1 + x|subject] + [1|trial]) with participants that replied True to Question 8 (Q8_T_; see **Table 2**) as reference group. ***A***, Illustration of the posterior predictive checks where the distribution of the observed outcome variable (y, in our case performance tempo) is compared to simulated datasets (y_rep_) from the posterior predictive distribution (100 draws). ***B***, Distribution of the difference in ms between performance tempo (intercept) in Q8_T_ and in participants that replied False to Question 8 (Q8_F_; see **Table 2**). The grey vertical bar indicates the posterior point estimate, while the grey area under the curve represents the 95% credible interval (CI). In the current plot, CI does overlap with zero (the null hypothesis). This indicates that there is 95% probability of no between-group differences in performance tempo. ***C***, Results of the BLMM analysis. We analysed how the strength of predictions about the action-reward contingency modulates performance tempo separately for Q8_T_ (in dark brown) and Q8_F_ (in light brown). Here, mIKI (performance tempo: mean inter-keystroke-interval) values are represented in the log-scale. The negative slopes suggest that stronger predictions about the action-reward contingency are associated with faster performance tempo, which replicates our findings in the main experiment (see Figure 5C). ***D***, Distribution of the difference between slopes in Q8_T_ and Q8_F_. Here, as CIs include zero we can conclude with 95% probability that groups do not differ in how the strength of predictions about the reward contingency influences motor performance tempo. Thus, the sensitivity of performance tempo to the strength of predictions about the reward mapping is not differently modulated between groups.

**Table 9.** Assessment of predictive accuracy in the models on performance tempo used in the control experiment.

| Model # | ELPD_difference | SE_difference |
| --- | --- | --- |
| 6 | 0 | 0 |
| 5 | -144.2029 | 20.33526 |
| 4 | -215.8902 | 24.35042 |
| 3 | -218.3484 | 24.53194 |
| 2 | -293.3391 | 29.49327 |
| 1 | -293.4484 | 29.49365 |

Overall, we found no differences between Q8_T_ and Q8_F_ in performance tempo, which is consistent with our previous between-group analyses in the control sample. The posterior estimate of the intercept in Q8_T_ (the reference group) was equal to 5.62, CI = [5.51, 5.73] (in ms: 275, CI [246, 307]). The distribution of the difference between intercepts in Q8_T_ and Q8_F_ had a posterior estimated value of 0.07, CI = [-0.12, 0.26] (in ms, 20, CI [-34, 75) (**Figure 7B**). Zero was included in the 95% CI of the latter distribution, suggesting that subjective inferences about credit assignment did not impact performance tempo.

We found a negative association (slope) between the strength of predictions about the action-reward contingency and performance tempo. This replicates our main findings showing that stronger predictions about the reward contingencies are followed by faster execution tempo (**Figure 7B**). Specifically, the 95% CI of the posterior distribution for the fixed effect of x (slope of the association between y and x in the reference group, Q8_T_) did not include zero, with a posterior estimate of -0.04, CI [-0.06, -0.01].

Further, no between-group slope differences were observed. Indeed, the distribution of the interaction effect group * x included zero (posterior estimate = 0, CI = [-0.04, 0.04]). Thus, subjective inferences about the causes for the absence of reward did not modulate the sensitivity of performance tempo to the strength of expectations about the action-reward contingency (**Figure 7D**).

In addition, the estimates of the standard deviation for the random effects by subject of the intercept (measure of inter-individual variability in performance tempo intercepts) and x (index of slope differences between individuals) were 0.27, CI = [0.21, 0.35] and 0.06, CI = [0.04, 0.08], respectively. The estimate of the standard deviation for the random effect of the intercept by trial was 0.08, CI = [0.06, 0.09], informing about the trial-by-trial variability in performance tempo. See **Table 10** for a summary of the posterior distributions for the winning model.

**Table 10.** Summary of the posterior distributions for the fixed effects of model number 6 on performance tempo in the control experiment.

| Fixed Effect | Estimate | l-95% CI | u-95% CI | R-hat |
| --- | --- | --- | --- | --- |
| y: Q8<sub>T</sub> | 5.62 | 5.51 | 5.73 | 1.00 |
| y: Q8<sub>T</sub> vs Q8<sub>F</sub> | 0.07 | -0.12 | 0.26 | 1.00 |
| x: Q8<sub>T</sub> | -0.04 | -0.06 | -0.01 | 1.00 |
| group * x: Q8<sub>T</sub> vs Q8<sub>F</sub> | -0.00 | -0.04 | 0.04 | 1.00 |

Estimates, credible intervals (CIs) and R-hat values for the fixed effects of the best fitting model in the control study (model number 6, $y \sim 1 + \text{group} * x + [1 + x|\text{subject}] + [1|\text{trial}]$ ) in brms with performance tempo as dependent variable. y: Q8<sub>T</sub> refers to the posterior estimate for the intercept in the reference group (participants that replied True to Question 8, Q8<sub>T</sub>). y: Q8<sub>T</sub> vs Q8<sub>F</sub> reflects the posterior distribution of the difference between intercepts (Q8<sub>T</sub> vs participants that replied False to Question 8 [Q8<sub>F</sub>]) x: Q8<sub>T</sub> is the posterior distribution of the association (slope) between performance tempo and the strength of predictions about the action-reward

Finally, we investigated the effect of differences in inferences about reward assignment on post-performance subjective error rate. First, the subjective error rate estimation was validated by computing BF analysis on the correlation between subjective and empirical error rates. Results provided strong evidence for a positive association in the full sample (N = 39; BF = 10.204; r = 0.448, p = 0.004). Next, we found no support for between-group differences in the subjective error rate (BF = 0.432, demonstrating anecdotal evidence for the null hypothesis; t_(36)_ = -0.850, p = 0.401). Thus, being not *always* sure about the causes for the lack of reward did not influence the rate of subjective number estimate of performance errors.

To conclude, our analyses provided evidence for the lack of differences between Q8_T_ and Q8_F_ in the evaluated parameters, suggesting that subjective inferences about task-related reward assignment do not modulate decision-making, general motor performance or the association between expectation on reward probability and motor vigour. Thus, whether groups in our main experiment had differences in credit assignment or not would have likely not led to a modulation of group effects. In addition, here we found further support for our main research hypothesis, whereby stronger predictions about the action-reward contingency enhanced motor vigour through faster performance.

## DISCUSSION

We investigated how predictions about the tendency of the action-reward contingency invigorated motor performance trial-by-trial in healthy younger adults (HYA), in medicated Parkinson’s Disease patients (PD), and in an age-matched sample of healthy older adults (HOA). We used a novel combination of a standard one-armed bandit decision-making paradigm and a motor sequence task. We then fitted the trial-by-trial behavioural data using the Hierarchical Gaussian Filter (HGF; Mathys et al. 2011, 2014; Frässle et al., 2021) and performed Bayesian analyses (Bayes Factor and Bayesian Linear Mixed Models [BLMM]). The results demonstrated a trial-by-trial modulation of performance tempo by the strength of expectations on the action-reward contingencies. Moreover, BLMM revealed a similar sensitivity of performance tempo to these predictions in HYA, HOA and PD. PD and HOA were, however, generally slower. This invigoration effect was limited to performance tempo, as the strength of predictions about the reward contingencies did not modulate reaction times (RT).

Our findings expand on previous investigations of the beneficial effects of reward on motor behaviour (e.g., faster and more accurate motor performance, increased velocity; Sedaghat- Nekad et al., 2019), which have been limited to manipulations of reward magnitude (presence/absence; large/small) in deterministic contexts (Codol et al., 2020; Sporn et al., 2020; Adkins & Lee, 2021; Aves et al., 2021). Here we showed that dynamically updating beliefs about the reward probabilities influenced performance tempo trial-by-trial. This converges with computational work demonstrating that updating of beliefs in a perceptual task speeds RT (Marshall et al., 2016). The authors found that, as participants learned to track the transition probabilities between stimuli, the model estimates of different decision-making variables affected RT. We revealed that the trial-by-trial influence of motor vigour by belief updating can be extended beyond the perceptual domain to learning about action-reward contingencies. Marshall et al. (2016) further showed that dopaminergic (DA) receptor antagonism attenuated the sensitivity of RT to uncertainty estimates. Here, our BLMM analyses demonstrated in HYA, HOA and PD a similar sensitivity (slope) of performance tempo to the strength of predictions about the reward mapping. Hence, the positive effect of updating beliefs about the changing relationships between actions and the corresponding outcomes on motor performance did not deteriorate with age, or in medicated PD. This is consistent with evidence pointing towards preserved reward sensitivity and probabilistic learning in ageing and medicated PD (Fera et al., 2005; Euteneuer et al., 2009; Pietschmann et al., 2011; Gescheidt et al., 2012, 2013; Chen et al., 2020; Aves et al., 2021). Yet, these results have been challenged by other work showing impaired decision making and reward-based learning in both groups (Cools et al., 2001; Shohamy et al., 2004; Weiler et al., 2008 Eppinger et al., 2011; Nassar et al., 2016; Hämmerer et al., 2019). Moreover, there was a lack of evidence on how reward sensitivity and reversal learning interact to modulate motor vigour in healthy ageing and PD. In our study, we provided compelling evidence for a preservation of motor invigoration by expectations of reward probability in HOA and PD.

Surprisingly, we found no modulation of RT by the strength of predictions about the action-reward contingencies. The sensitivity of RT to reward has been assessed in perceptual and decision-making tasks, demonstrating reduced RT as a function of reward expectation (Guitart-Masip et al., 2011; Milstein & Dorris, 2011; Marshall et al., 2016 Griffiths & Beierholm, 2017). Evidence in the field of motor control has also shown smaller RT for movement paired with rewards (Summerside et al., 2018). However, overall little attention has been given to reward-driven modulation of RT in motor tasks (e.g., focus on movement times, rather than RT [Sporn et al., 2020; Aves et al., 2021]). In our study, inferences about the probabilistic structure of the reward contingencies invigorated performance tempo trial-by-trial without affecting RT. A speculative interpretation is that not prompting participants to complete the trial as fast as possible may have introduced extra variability in RT. Follow-up work using different task instructions could help explain our results.

We found extreme evidence for between-group differences in the mean performance tempo. Indeed, HYA were faster than HOA and PD, and HOA quicker than PD. The slower sequence execution in HOA is consistent with a general slowness of hand movements in later stages of life (Goggin & Meeuwsen, 1992; Ketcham et al., 2002; Aves et al., 2021). Regarding PD, the slower performance is likely explained by a sequence effect (SE). SE is a common bradykinetic symptom in PD, which manifests through a slower and decreased in amplitude execution of sequential movements (Kang et al., 2010). DA intake does not ameliorate symptoms associated with SE, suggesting a non-DA involvement in the pathophysiology of this effect (Kang et al., 2010; Bologna et al., 2016). Similar results were found for RT, with HYA displaying shorter RT than HOA and PD. Yet, contrary to performance tempo, RT did not dissociate between HOA and PD.

We additionally found evidence for similar win and error rates in our three groups. This converges with research arguing for preserved probabilistic decision making in medicated PD (Euteneuer et al., 2009; Gescheidt et al., 2012, 2013). Yet, groups differed in the number of sequence renditions during familiarisation, with HOA and PD practising more compared to HYA. Thus, the ability to complete our task was not compromised with age nor in PD, despite the latter requiring extra practice to initially memorise the sequences and displaying a general movement slowness.

Groups also did not differ in the main markers of decision making. We provided some evidence for the absence of a group effect on tonic volatility (*ω*_2_), an index of individual learning about the action-reward mapping under volatility (Hein et al., 2021). Similarly, the estimated uncertainty about the action-reward tendency (*σ*_2_) did not differ between groups. Finally, groups also exhibited a similar mapping from beliefs to responses, driven by the response model parameter ζ. A limitation is that numerical instability prevented us from including the HGF_3_ in the model comparison, although qualitative evidence in 80% of the participants in which the HGF_3_ converged supported that the HGF_2_ and HGF_3_ had very similar LME values (differences < 1). On the one hand, the absence of group effects on decision making is consistent with computational work suggesting that learning about the probabilistic task structure is independent from DA neurotransmission (Marshall et al., 2016). Specifically, the authors found that DA antagonism did not affect beliefs updating about task volatility and transition probability between stimuli. On the other hand, neuroimaging evidence suggests that probabilistic learning relies on DA signalling in mesolimbic and prefrontal circuits (de Boer et al., 2017). In this scenario, drug intake might have restored depleted DA levels in our medicated PD sample, therefore allowing for the dynamic update of beliefs about the reward probabilities. This, however, contradicts previous studies arguing for impaired reversal learning and probabilistic decision making in medicated PD (Cools et al., 2006; Gotham et al., 1998; Vaillancourt et al., 2013). These findings have been contextualised within the “overdose hypothesis” framework according to which DA intake acts by restoring depleted DA in the motor dorsal striatal pathway, yet overdosing the relatively spared ventral portion (supporting probabilistic and reversal learning) (Ferrazzoli et al., 2016; Torta & Castelli, 2008). Overall, the effects of DA-replacement therapy on decision making are still poorly understood (Ryterska et al., 2013; Kjær et al., 2019).

Our main study did not determine whether participants correctly inferred the hidden causes for the lack of reward (McDougle et al., 2016). Our control experiment demonstrated that, when participants can be dissociated in their tendency for credit assignment, this does not result in differences in general motor performance, decision making or motor vigour. Individual inferences about credit assignment did not explain differences in the subjective number estimate of performance errors either. Because the feedback that participants received was veridical (unlike in McDougle et al., 2016), the effects of misattribution of the causes of zero reward in our study are likely very small, as the anecdotal evidence suggests. Moreover, as most participants reported using sound and finger movements to detect errors, the misattribution – even when it happened – was infrequent.

A potential confounding factor in our main results could be related to PD not being monetarily rewarded. Nonetheless, PD showed similar decision making and learning motifs to paid participants, which makes a role of money in accounting for our results unlikely. Finally, as some participants completed the task in Italian, we controlled for the role of language. We showed converging outcomes when contrasting PD to either the full HOA sample or to the Italian HOA subsample. Hence, between-group effects on general task performance and decision making cannot be accounted for by language differences.

To conclude, this study is the first to demonstrate that inferring the probabilistic reward mappings positively biases motor performance through faster execution tempo. Additionally, we provided novel evidence for a preserved sensitivity of the motor invigoration effects in HOA and PD. Thus, healthy young, old and medicated PD can similarly obtain benefits in their motor performance when updating beliefs about the volatile action-reward contingencies.

## DATA AVAILABILITY

The data that support some of the findings of this study are available from the Open Science Framework Data Repository under the accession code a9wbu. https://osf.io/7kfbj/?view_only=6e5ed0c8216f48b3964b1f9df117fa36

## CODE AVAILABILITY

Code for the main brms analysis in the first experiment has been deposited in the Open Science Framework Data Repository under the accession code a9wbu. https://osf.io/7kfbj/?view_only=6e5ed0c8216f48b3964b1f9df117fa36

## Acknowledgments

We would like to thank Osama Shah for programming the task in JavaScript

## REFERENCES

Adkins TJ, Lee TG (2021) Reward modulates cortical representations of action. NeuroImage 228:117708.

Andraszewicz S, Scheibehenne B, Rieskamp J, Grasman R, Verhagen J, Wagenmakers E-J (2015) An Introduction to Bayesian Hypothesis Testing for Management Research. J Manag 41:521–543.

Aves P, Moreau L, Alghamdi A, Sporn S, Galea JM (2021) Age-Related Differences in Reward-Based Modulation of Sequential Reaching Performance. bioRxiv 461920. https://doi.org/10.1101/2021.09.27.461920;

Behrens TEJ, Woolrich MW, Walton ME, Rushworth MFS (2007) Learning the value of information in an uncertain world. Nat Neurosci 10:1214–1221.

Bologna M, Leodori G, Stirpe P, Paparella G, Colella D, Belvisi D, Fasano A, Fabbrini G, Berardelli A (2016) Bradykinesia in early and advanced Parkinson’s disease. J Neurol Sci 369:286–291.

Botvinick M, Braver T (2015) Motivation and Cognitive Control: From Behavior to Neural Mechanism. Annu Rev Psychol 66:83–113.

Bürkner P-C (2017) brms: An *R* Package for Bayesian Multilevel Models Using *Stan*. J Stat Softw 80:1–28.

Bürkner P-C (2018) Advanced Bayesian Multilevel Modeling with the R Package brms. R J 10:395–411.

Bürkner P-C (2021) Bayesian Item Response Modeling in R with brms and Stan. J Stat Softw 100:1–54.

Carroll TJ, McNamee D, Ingram JN, Wolpert DM (2019) Rapid Visuomotor Responses Reflect Value-Based Decisions. J Neurosci 39:3906–3920.

Chen X, Voets S, Jenkinson N, Galea JM (2020) Dopamine-Dependent Loss Aversion during Effort-Based Decision-Making. J Neurosci 40:661–670.

Codol O, Holland PJ, Manohar SG, Galea JM (2020) Reward-Based Improvements in Motor Control Are Driven by Multiple Error-Reducing Mechanisms. 40: 3604–3620.

Cools R (2001) Enhanced or Impaired Cognitive Function in Parkinson’s Disease as a Function of Dopaminergic Medication and Task Demands. Cereb Cortex 11:1136–1143.

Cools R (2006) Dopaminergic modulation of cognitive function-implications for l-DOPA treatment in Parkinson’s disease. Neurosci Biobehav Rev 30:1–23.

Cousineau D (2020) How many decimals? Rounding descriptive and inferential statistics based on measurement precision. J Math Psychol 97:102362.

de Berker AO, Rutledge RB, Mathys C, Marshall L, Cross GF, Dolan RJ, Bestmann S (2016) Computations of uncertainty mediate acute stress responses in humans. Nat Commun 7:10996.

de Boer L, Axelsson J, Riklund K, Nyberg L, Dayan P, Bäckman L, Guitart-Masip M (2017) Attenuation of dopamine-modulated prefrontal value signals underlies probabilistic reward learning deficits in old age. eLife 6:e26424.

den Ouden HEM, Daunizeau J, Roiser J, Friston KJ, Stephan KE (2010) Striatal Prediction Error Modulates Cortical Coupling. J Neurosci 30:3210–3219.

Diaconescu AO, Mathys C, Weber LAE, Daunizeau J, Kasper L, Lomakina EI, Fehr E, Stephan KE (2014) Inferring on the Intentions of Others by Hierarchical Bayesian Learning. PLoS Comput Biol 10:e1003810.

Eppinger B, Hämmerer D, Li S-C (2011) Neuromodulation of reward-based learning and decision making in human aging: Eppinger et al. Ann N Y Acad Sci 1235:1– 17.

Euteneuer F, Schaefer F, Stuermer R, Boucsein W, Timmermann L, Barbe MT, Ebersbach G, Otto J, Kessler J, Kalbe E (2009) Dissociation of decision-making under ambiguity and decision-making under risk in patients with Parkinson’s disease: A neuropsychological and psychophysiological study. Neuropsychologia 47:2882–2890.

Fahn S, Elton RL (1987) The unified Parkinson’s disease rating scale. In: Recent developments in Parkinson’s disease, Vol 2 (Fahn S, Marsden CD, Calne DB, Goldstein M, eds), pp 153–163, 293–304. Florham Park, NJ. Macmillan Health Care Information.

Feldman H, Friston KJ (2010) Attention, Uncertainty, and Free-Energy. Front Hum Neurosci 4:215.

Fera F (2005) Neural Mechanisms Underlying Probabilistic Category Learning in Normal Aging. J Neurosci 25:11340–11348.

Ferrazzoli D, Carter A, Ustun FS, Palamara G, Ortelli P, Maestri R, Yücel M, Frazzitta G (2016) Dopamine Replacement Therapy, Learning and Reward Prediction in Parkinson’s Disease: Implications for Rehabilitation. Front Behav Neurosci 10:121.

Frässle S et al. (2021) TAPAS: An Open-Source Software Package for Translational Neuromodeling and Computational Psychiatry. Front Psychiatry 12:68081.

Friston KJ, Stephan KE, Montague R, Dolan RJ (2014) Computational psychiatry: the brain as a phantastic organ. Lancet Psychiatry 1:148–158.

Galaro JK, Celnik P, Chib VS (2019) Motor Cortex Excitability Reflects the Subjective Value of Reward and Mediates Its Effects on Incentive-Motivated Performance. J Neurosci 39:1236–1248.

Gelman A, Rubin DB (1992) Inference from Iterative Simulation Using Multiple Sequences. Stat Sci 7:457–472.

Gescheidt T, Czekóová K, Urbánek T, Mareček R, Mikl M, Kubíková R, Telecká S, Andrlová H, Husárová I, Bareš M (2012) Iowa Gambling Task in patients with early-onset Parkinson’s disease: strategy analysis. Neurol Sci 33:1329–1335.

Gescheidt T, Mareček R, Mikl M, Czekóová K, Urbánek T, Vaníček J, Shaw DJ, Bareš M (2013) Functional anatomy of outcome evaluation during Iowa Gambling Task performance in patients with Parkinson’s disease: an fMRI study. Neurol Sci 34:2159–2166.

Goggin NL, Meeuwsen HJ (1992) Age-related differences in the control of spatial aiming movements. Res Q Exerc Sport 63:366–372.

Gotham AM, Brown RG, Marsden CD (1988) ‘Frontal’ cognitive function in patients with Parkinson’s disease ‘on’ and ‘off’ levodopa. Brain 111:299–321.

Griffiths B, Beierholm UR (2017) Opposing effects of reward and punishment on human vigor. Sci Rep 7:42287.

Guitart-Masip M, Beierholm UR, Dolan R, Duzel E, Dayan P (2011) Vigor in the Face of Fluctuating Rates of Reward: An Experimental Examination. J Cogn Neurosci 23:3933–3938.

Hämmerer D, Schwartenbeck P, Gallagher M, FitzGerald THB, Düzel E, Dolan RJ (2019) Older adults fail to form stable task representations during model-based reversal inference. Neurobiol Aging 74:90–100.

Hein TP, de Fockert J, Ruiz MH (2021) State anxiety biases estimates of uncertainty and impairs reward learning in volatile environments. NeuroImage 224:117424.

Hein TP, Herrojo Ruiz M (2022) State anxiety alters the neural oscillatory correlates of predictions and prediction errors during reward-based learning. NeuroImage 249:118895.

Herrojo Ruiz M, Jabusch H-C, Altenmüller E (2009) Detecting Wrong Notes in Advance: Neuronal Correlates of Error Monitoring in Pianists. Cereb Cortex 19:2625–2639.

Herrojo Ruiz M, Maess B, Altenmüller E, Curio G, Nikulin VV (2017) Cingulate and cerebellar beta oscillations are engaged in the acquisition of auditory-motor sequences. Hum Brain Mapp 38:5161–5179.

Iglesias S, Mathys C, Brodersen KH, Kasper L, Piccirelli M, den Ouden HEM, Stephan KE (2013) Hierarchical Prediction Errors in Midbrain and Basal Forebrain during Sensory Learning. Neuron 80:519–530.

Kang SY, Wasaka T, Shamim EA, Auh S, Ueki Y, Lopez GJ, Kida T, Jin S-H, Dang N, Hallett M (2010) Characteristics of the sequence effect in Parkinson’s disease: Sequence Effect in Parkinson’s Disease. Mov Disord 25:2148–2155.

Ketcham CJ, Seidler RD, Van Gemmert AWA, Stelmach GE (2002) Age-Related Kinematic Differences as Influenced by Task Difficulty, Target Size, and Movement Amplitude. J Gerontol B Psychol Sci Soc Sci 57:P54–P64.

Kjær SW, Damholdt MF, Callesen MB (2018) A systematic review of decision-making impairments in Parkinson’s Disease: Dopaminergic medication and methodological variability. Basal Ganglia 14:31–40.

Lewandowski D, Kurowicka D, Joe H (2009) Generating random correlation matrices based on vines and extended onion method. J Multivar Anal 100:1989–2001.

Manohar SG, Muhammed K, Fallon SJ, Husain M (2019) Motivation dynamically increases noise resistance by internal feedback during movement. Neuropsychologia 123:19–29.

Marshall L, Mathys C, Ruge D, de Berker AO, Dayan P, Stephan KE, Bestmann S (2016) Pharmacological Fingerprints of Contextual Uncertainty. PLOS Biol 14:e1002575.

Mathys C (2011) A Bayesian foundation for individual learning under uncertainty. Front Hum Neurosci 5:1–20.

Mathys CD, Lomakina EI, Daunizeau J, Iglesias S, Brodersen KH, Friston KJ, Stephan KE (2014) Uncertainty in perception and the Hierarchical Gaussian Filter. Front Hum Neurosci 8:1–24.

MATLAB and Statistics Toolbox Release 2020b, The MathWorks, Inc., Natick, Massachusetts, United States.

McDougle SD, Boggess MJ, Crossley MJ, Parvin D, Ivry RB, Taylor JA (2016) Credit assignment in movement-dependent reinforcement learning. Proc Natl Acad Sci 113:6797–6802.

Metitieri T, Geroldi C, Pezzini A, Frisoni GB, Bianchetti A, Trabucchi M (2001). The Itel-MMSE: An Italian telephone version of the mini-mental state examination. Int J Geriatr Psychiatry 16:166–167.

Milstein DM, Dorris MC (2011) The Relationship between Saccadic Choice and Reaction Times with Manipulations of Target Value. Front Neurosci 5:122.

Nassar MR, Bruckner R, Gold JI, Li S-C, Heekeren HR, Eppinger B (2016) Age differences in learning emerge from an insufficient representation of uncertainty in older adults. Nat Commun 7:11609.

Padmala S, Pessoa L (2011) Reward Reduces Conflict by Enhancing Attentional Control and Biasing Visual Cortical Processing. J Cogn Neurosci 23:3419– 3432.

Pietschmann M, Endrass T, Czerwon B, Kathmann N (2011) Aging, probabilistic learning and performance monitoring. Biol Psychol 86:74–82.

Powers AR, Mathys C, Corlett PR (2017) Pavlovian conditioning–induced hallucinations result from overweighting of perceptual priors. Science 357:596– 600.

R Core Team (2022). R: A language and environment for statistical computing. R Foundation for Statistical Computing, Vienna, Austria. URL https://www.R-project.org/.

Reed EJ, Uddenberg S, Suthaharan P, Mathys CD, Taylor JR, Groman SM, Corlett PR (2020) Paranoia as a deficit in non-social belief updating. eLife 9:e56345.

Rescorla RA, Wagner AR (1972) A theory of Pavlovian conditioning: variations in the effectiveness of reinforcement and nonreinforcement. Class. Cond. II: Curr. Res. Theory 2:64–99.

Rouder JN, Morey RD, Speckman PL, Province JM (2012) Default Bayes factors for ANOVA designs. J Math Psychol 56:356–374.

Ryterska A, Jahanshahi M, Osman M (2013) What are people with Parkinson’s disease really impaired on when it comes to making decisions? A meta-analysis of the evidence. Neurosci Biobehav Rev 37:2836–2846.

Sedaghat-Nejad E, Herzfeld DJ, Shadmehr R (2019) Reward Prediction Error Modulates Saccade Vigor. J Neurosci 39:5010–5017.

Sheffield JM, Suthaharan P, Leptourgos P, Corlett PR (2022) Belief Updating and Paranoia in Individuals with Schizophrenia. Biol Psychiatry Cogn Neurosci Neuroimaging doi: https://doi.org/10.1016/j.bpsc.2022.03.013.

Shohamy D (2004) Cortico-striatal contributions to feedback-based learning: converging data from neuroimaging and neuropsychology. Brain 127:851–859.

Sivula T, Magnusson M, Matamoros AA, Vehtari A (2022) Uncertainty in Bayesian Leave-One-Out Cross-Validation Based Model Comparison. arXiv 2008.10296. https://doi.org/10.48550/arXiv.2008.10296.

Soch J, Allefeld C (2018) MACS – a new SPM toolbox for model assessment, comparison and selection. J Neurosci Methods 306:19–31.

Spielberger CD, Gorsuch RL, Lushene R, Vagg PR, Jacobs GA (1983). Manual for the State-Trait Anxiety Inventory. Palo Alto, CA: Consulting Psychologists Press.

Sporn S, Chen X, Galea JM (2020) Reward-based invigoration of sequential reaching. bioRxiv 152876. https://doi.org/10.1101/2020.06.15.152876

Stefanics G, Heinzle J, Horváth AA, Stephan KE (2018) Visual Mismatch and Predictive Coding: A Computational Single-Trial ERP Study. J Neurosci 38:4020–4030.

Summerside EM, Shadmehr R, Ahmed AA (2018) Vigor of reaching movements: reward discounts the cost of effort. J Neurophysiol 119:2347–2357.

Sutton RS (1992) Gain adaptation beats least squares? Paper presented at the 7th Yale Workshop on Adaptive and Learning Systems, New Haven, CT, May.

Torta DME, Castelli L (2008) Reward pathways in Parkinson’s disease: Clinical and theoretical implications. Psychiatry Clin Neurosci 62:203–213.

Vaillancourt DE, Schonfeld D, Kwak Y, Bohnen NI, Seidler R (2013) Dopamine overdose hypothesis: Evidence and clinical implications: Dopamine Overdose Hypothesis. Mov Disord 28:1920–1929.

Valton V, Wise T, Robinson OJ (2020) Recommendations for Bayesian hierarchical model specifications for case-control studies in mental health. arXiv 2011.01725. https://doi.org/10.48550/arXiv.2011.01725.

Vehtari A, Gelman A, Gabry J (2017) Practical Bayesian model evaluation using leave-one-out cross-validation and WAIC. Stat Comput 27:1413–1432.

Weber LA, Diaconescu AO, Mathys C, Schmidt A, Kometer M, Vollenweider F, Stephan KE (2020) Ketamine Affects Prediction Errors about Statistical Regularities: A Computational Single-Trial Analysis of the Mismatch Negativity. J Neurosci 40: 5658–5668.

Weiler JA, Bellebaum C, Daum I (2008) Aging affects acquisition and reversal of reward-based associative learning. Learn Mem 15:190–197.

Zigmong AS, Snaith RP (1983) The hospital anxiety and depression scale. Acta Psychiatr Scand 67:361–370.

